# Generative modeling of brain maps with spatial autocorrelation

**DOI:** 10.1101/2020.02.18.955054

**Authors:** Joshua B. Burt, Markus Helmer, Maxwell Shinn, Alan Anticevic, John D. Murray

## Abstract

Studies of large-scale brain organization have revealed interesting relationships between spatial gradients in brain maps across multiple modalities. Evaluating the significance of these findings requires establishing statistical expectations under a null hypothesis of interest. Through generative modeling of synthetic data that instantiate a specific null hypothesis, quantitative benchmarks can be derived for arbitrarily complex statistical measures. Here, we present a generative null model, provided as an open-access software platform, that generates surrogate maps with spatial autocorrelation (SA) matched to SA of a target brain map. SA is a prominent and ubiquitous property of brain maps that violates assumptions of independence in conventional statistical tests. Our method can simulate surrogate brain maps, constrained by empirical data, that preserve the SA of cortical, subcortical, parcellated, and dense brain maps. We characterize how SA impacts *p*-values in pairwise brain map comparisons. Furthermore, we demonstrate how SA-preserving surrogate maps can be used in gene ontology enrichment analyses to test hypotheses of interest related to brain map topography. Our findings demonstrate the utility of SA-preserving surrogate maps for hypothesis testing in complex statistical analyses, and underscore the need to disambiguate meaningful relationships from chance associations in studies of large-scale brain organization.

## Introduction

Recent technological advancements in neuroimaging, large-scale connectomics, and high-throughput transcriptomics have facilitated the discovery of conserved principles of brain organization (Huntenburg et al., 2018; Burt et al., 2018; Fornito et al., 2019). Studies of spatial representations of brain features — i.e., brain maps — have revealed large-scale gradients of microscale and macroscale features (Wagstyl et al., 2015; Margulies et al., 2016; Burt et al., 2018; Preller et al., 2018; Vázquez-Rodríguez et al., 2019; Fulcher et al., 2019; Royer et al., 2019). Furthermore, gradients from distinct modalities exhibit intriguing relationships, including topographic alignment of local cytoarchitectural variation (Wagstyl et al., 2015; Hilgetag et al., 2016), long-range connectivity (Markov et al., 2014), gene expression profiles (Burt et al., 2018; Anderson et al., 2018), neurophysiological properties (Murray et al., 2014), and participation in hierarchies of functionally specialized networks (Margulies et al., 2016; Burt et al., 2018; Wang, 2020). However, interpretation of statistical measures derived from brain maps requires the establishment of statistical expectations under a well-defined null hypothesis of interest.

We designate a null model as generative if it generates surrogate data which instantiate a specific null hypothesis (Fornito et al., 2016; Betzel and Bassett, 2017). Generative null modeling is particularly advantageous because surrogate data can be directly operated on to derive null distributions for arbitrarily complex statistical measures. For brain maps, randomly permuting values across regions as part of a permutation test can be considered a distribution-preserving generative null model which produces surrogate maps with randomized topographies. Yet the assumptions built into the null hypotheses of permutation testing, as well as conventional parametric testing, are strongly violated by a characteristic and ubiquitous property of brain maps: spatial autocorrelation (SA). Due to SA, values of brain features in spatially proximal regions tend to be more similar than values of spatially distant regions. Thus, statistical claims about a particular brain map topography should be evaluated against a generative null model in which that target map’s SA structure is explicitly incorporated into the null hypothesis.

There is growing appreciation in the neuroimaging field that innovative methods are required to account for the impact of SA on statistical analyses of large-scale brain maps and spatial gradients (Alexander-Bloch et al., 2018; Burt et al., 2018; de Wael et al., 2019). Recent proposals have focused primarily on deriving corrected *p*-values for tests of spatial correspondence between pairs of brain maps. The most widely adopted of these approaches, the spin test, involves randomizing the anatomical alignment between two cortical surface maps through spherical rotation by a random angle (Alexander-Bloch et al., 2018). However, the utility of this non-generative approach is limited, because it does not produce surrogate maps with complete cortical coverage, nor does it generalize to volumetric data. There remains a significant unmet methodological need for SA-preserving generative null modeling in neuroscience.

Here, we adapt a method from geostatistics to develop a generative null modeling framework for generating surrogate brain maps with SA matched to the SA of a target brain map (Viladomat et al., 2014). We first demonstrate that this method can be used to correct for the impact of SA on statistical significance values derived from pairwise brain map comparisons. After describing the statistical foundations of the model, we provide three illustrative applications to empirical data, contrasting model-derived results with results from conventional, spatially naive statistical tests. Our approach can be flexibly applied to a range of brain map representations, including surface-based and volumetric geometries at either parcellated or dense resolutions. We apply our method to gene ontology (GO) enrichment analyses, which are commonly used to infer biological correlates of brain map topographies. We found that GO results are spuriously driven by SA, and we develop a workflow for evaluating the GO enrichment of a map’s topography while controlling for its SA. We have developed a Python-based implementation of our method, with additional neuroimaging-specific functionality, which is released as an open-source software package for the field.

## Methods

The method described below is implemented in an open-access, Python-based software package, BrainSMASH: Brain Surrogate Maps with Autocorrelated Spatial Heterogeneity (https://github.com/murraylab/brainsmash).

### Generating spatial autocorrelation-preserving surrogate maps

We present the algorithm first proposed by Viladomat et al. (2014) for testing correlations between autocorrelated fields, here used to generate SA-preserving brain maps. The algorithm can be conceptually subdivided into two main steps:

1. Randomly permute the values in a target brain map.
2. Smooth and rescale the permuted map to recover lost SA structure.

Let x be a brain map whose value in brain region *i* is denoted *x_i_*. We randomly shuffle (i.e., permute) the values in x to obtain the permuted map 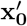. We perform a local kernel-weighted sum of values in 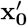 to construct the smoothed map 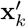, where the *i*-th element is computed as:

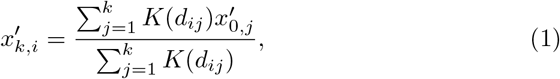

where *k* is the number of nearest neighboring regions used to perform the smoothing, *K* is a distance-dependent smoothing kernel, and *d_ij_* is the distance separating regions *i* and *j*. We use an exponentially decaying smoothing kernel with a characteristic length scale equal to the distance of the *k*-th nearest neighbor. Following Viladomat et al. (2014), our smoothing kernel is truncated, here at the characteristic length scale where it has a value of *e*^-1^. The distance at which the kernel truncates will therefore be larger in regions where the brain map is more sparsely sampled. The parameter *k*, which sets the spatial scale of the SA reintroduced into the surrogate map, is chosen from a set of user-defined values such that surrogate maps’ fit to the target map is maximized (which we will return to below).

After smoothing the permuted map, 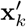 must be rescaled such that its SA approximately matches the SA in the target map. To do this, we construct a variogram—a summary measure of the autocorrelation in spatial data—for each brain map. The variogram, which provides a measure of pairwise variation as a function of distance, is typically computed within finite-width distance intervals: in the distance interval centered around length scale *h* with width 2*δ*, the value of the variogram, denoted *γ*, for brain map x is equal to the sample variance:

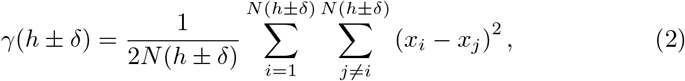

where *N* (*h* ± *δ*) is the number of sample pairs separated by a distance *d_ij_* which lies in the interval *h* – *δ* ≤ *d_ij_* < *h* + *δ*.

Following Viladomat et al. (2014), we further reduce noise in the data by smoothing the variogram. To do this, we replace Equation 2 with

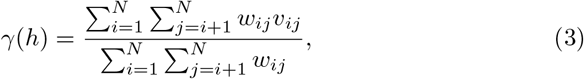

where 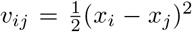, and the weights *w_ij_* are computed using a Gaussian kernel which falls off smoothly with distance:

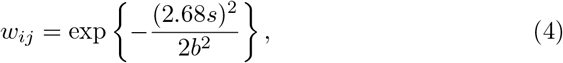

where *s* = ||*h* – *d_ij_*||, *d_ij_* is the distance between regions *i* and *j*, the bandwidth *b* controls the smoothness of the smoothed variogram, and constants are chosen such that the quartiles of the kernel are at ±0.25b (Viladomat et al., 2014). In other words, a pair of regions *i* and *j* contribute most strongly to the smoothed variogram evaluated at length scale *h* when their distance *d_ij_* is equal to *h*. In addition, because SA is primarily a local effect, only pairs of regions whose distance *d_ij_* lies in the bottom 25th percentile of the distribution {*d_ij_*} contribute to the weighted sum in Equation 3, following Viladomat et al. (2014). Throughout this study, Equation 3 was evaluated at 25 uniformly spaced distance intervals {*h*} across the range of distances *d_ij_* which fell in the bottom 25th percentile of all elements in the distance matrix **D**. The bandwidth b was chosen to be three times the distance interval spacing, i.e., *b* = 3 (*h_i_* – *h*_*i*-1_).

To recover SA in our surrogate maps, we maximize the fit between the target brain map’s variogram, *γ*[x], and the variogram of the smoothed map, 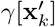, where *γ*[x] refers to Equation 3 evaluated at all {*h*} for brain map x. By matching the variogram of the smoothed map to the variogram of the target map, we impart the smoothed map with the characteristic SA structure of the target map. For each value of smoothing parameter *k*, we linearly regress 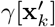 onto *γ*[x]; this procedure amounts to choosing a linearly transformed variogram from the family

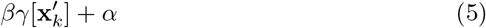

that is maximally similar to *γ*[x]. For each value of *k*, we compute the sum of squared errors (SSE) in the fit between *γ*[x] and the linearly transformed 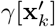. We then select the value of *k* which minimizes SSE, denoted *k*\*, to construct a surrogate map whose SA is approximately matched to the SA in *x*. The regression coefficients for *k*\*, denoted *α_k*_* and *β_k*_*, define the linear transformation of 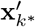 into the surrogate map 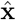:

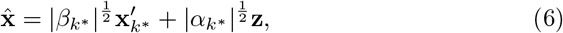

where z is a map of normally distributed random variates with zero mean and unit variance.

Constructing dense (i.e., vertex-or voxel-wise) surrogate brain maps imposes additional computational challenges. The number of elements in a distance matrix or variogram — that is, in a pairwise measure — scales like 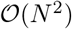, where *N* is the number of brain regions. Consequently, each pairwise measure for a dense map individually requires ~ 4GB of RAM, quickly exhausting the resources of standard laptop computers. To overcome this challenge, we developed an additional protocol for constructing dense surrogate maps. Because SA is primarily a local effect, only a subset of the smallest elements in a distance matrix, representing pairs of spatially proximal regions, are needed to construct reliable dense surrogate maps. For each row of a dense distance matrix, we therefore keep only the *k_nn_* smallest elements that are greater than zero. In other words, for each vertex/voxel, we keep only the distances to its knn nearest neighboring vertices/voxels. In this study, we used *k_nn_* = 1, 000 for dense cortical surrogate maps, and *k_nn_* = 1, 500 for cerebellar surrogate maps. In addition, following Viladomat et al. (2014), we approximated dense variograms using a random sampling of the data: for each dense surrogate map, we randomly sample *n_s_* regions (without replacement) to perform the variogram fitting procedure. We used *n_s_* = 1, 000 for dense cortical surrogates and *n_s_* = 500 for cerebellar surrogates. These two sampling techniques reduced the memory burden by a factor of ~ 900 for our dense cortical data, and result in a computational cost which scales linearly with the number of regions.

We have also developed an implementation which leverages memory-mapped arrays, such that distance matrices stored locally on disk are read into memory on an as-needed basis. The specific algorithms for constructing both dense and parcellated surrogate brain maps are provided in Appendix A. All key parameters described above are configurable in BrainSMASH and default to the values used in this study. More details can be found in the BrainSMASH documentation: https://brainsmash.readthedocs.io/.

### Data

#### Parcellated structural neuroimaging maps

Human T1w/T2w and cortical thickness maps in the surface-based CIFTI file format were obtained from the Human Connectome Project (HCP) (Van Essen et al., 2013). To produce the T1w/T2w maps, high resolution T1- and T2-weighted images were first registered to a standard reference space using an areal-feature-based technique (Glasser et al., 2016a; Robinson et al., 2014), then corrected for bias-field intensity inhomogeneities (Glasser and Van Essen, 2011; Glasser et al., 2013). Group-averaged (N = 339) left-hemispheric T1w/T2w and thickness maps were parcellated into 180 regions using the HCP’s Multi-Modal Parcellation (MMP1.0) (Glasser et al., 2016a). Assignment of MMP1.0 parcels to functional networks was performed through community detection analysis (Ito et al., 2017) on time-series correlations in the HCP resting-state fMRI dataset.

#### Gene expression maps

Gene expression data were pre-processed following a procedure which we previously reported (Burt et al., 2018). Briefly, we constructed gene expression maps using data from the Allen Human Brain Atlas (AHBA)—a publicly available transcriptional atlas of DNA microarray data, containing samples from hundreds of histologically validated neuroanatomical structures across six normal post-mortem human brains (Hawrylycz et al., 2012, 2015). Microarray expression data and all accompanying metadata were downloaded from the AHBA (http://human.brain-map.org). The raw microarray expression data for each of the six donors includes expression levels of 20,737 genes; our preprocessing pipeline yielded group-averaged gene expression profiles for 16,088 genes across 180 parcels in the left cortical hemisphere. Brain-specific genes were selected as in Burt et al. (2018).

#### Distance matrices

Matrices of three-dimensional Euclidean distance were used for subcortical analyses, while matrices of surface-based geodesic distance were used for cortical analyses. Geodesic distances between grayordinate vertices in the midthickness surface file were computed using the Connectome Workbench software. To compute the geodesic distance between two parcels *i* and *j*, we computed the average of all pairwise surface-based distances between a grayordinate vertex in parcel *i* and a vertex in parcel *j*.

### Gaussian random fields

To theoretically characterize the impact of SA, we simulated Gaussian random fields (GRFs) on a square lattice while parametrically varying SA. An important statistical feature of a random field is its autocorrelation function which, for isotropic and homogeneous random fields, is related to the power spectral density via the Wiener-Khinchin theorem. We therefore use a parametric function for the power spectral density to vary the SA of simulated GRFs:

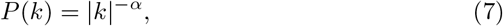

where *α* is a positive number and *k* is a spatial frequency (not to be confused with the number of nearest neighboring brain regions). We simulated GRFs on uniformly spaced two-dimensional grids within the unit interval with *N* tilings in each dimension. More details about the theoretical foundation of this approach are provided in Appendix B.

### Gene ontology enrichment analyses

Gene ontology (GO) enrichment analyses were scripted in the Python programming language using the GOATOOLS package (Klopfenstein et al., 2018). To generate gene sets used for these analyses, we performed partial least squares (PLS)-based cross-decomposition between 16, 088 gene expression maps and one brain map to identify genes whose spatial expression patterns were most strongly associated with a brain map’s topography (Vértes et al., 2016; Whitaker et al., 2016; Romero-Garcia et al., 2018; Morgan et al., 2019). For each brain map, we first identified the 1, 000 most strongly associated genes using PLS with a single latent variable, corresponding to the 1, 000 genes with largest positive PLS scores. For these genes, we then used GOATOOLS to identify significantly enriched annotations (i.e., GO categories) for these genes, as well as their Bonferroni-corrected significance values. All available UniProt IDs in the GOATOOLS database were used as the background reference set.

Prior to reporting enriched GO categories, following Vértes et al. (2016), we eliminated semantically redundant terms using the web-based tool REViGO (http://revigo.irb.hr) (Supek et al., 2011). To generate Fig. 7B, we first computed the number of surrogate brain maps which were enriched for each GO category. We kept the categories for which at least 5% of surrogate maps were significantly enriched. Our input to REViGO was this list of GO categories and the associated numbers of significantly enriched surrogate brain maps. In the web-based tool, we set the allowed similarity to “Medium (0.7)”, and we selected the option for numbers associated with each GO category to be “some other quantity, where higher is better.” Advanced options were left as their default values. REViGO-generated outputs for each of the three enrichment classes (biological process, molecular function, and cellular component) were exported as CSV files and aggregated. Finally, we applied thresholds of *u* > 0.9 for column “uniqueness”, *d* < 0.05 for column “dispensability”, and *v* > 60 for column “value”, which reduced the list to a set of 12 highly enriched and semantically unique categories that were subsequently plotted in semantic space.

### Principal components analysis

For a set of *N* genes, each with group-averaged gene expression values in *p* cortical parcels, we constructed a gene expression matrix *G* with one row for each cortical parcel and one column for each unique gene (i.e., with dimensions *P* × *N*). The *P* × *P* spatial covariance matrix *C* was constructed by computing the covariance between vectors of gene expression values for each pair of cortical parcels: *C_ij_* = Cov(*G_i_, G_j_*), where *G_i_* is the *i*th row in the matrix *G*, corresponding to the vector of *N* gene expression values for the *i*th cortical parcel. Eigendecomposition was performed on the spatial covariance matrix to obtain the matrix eigenvectors (i.e., the principal components, PCs) and their corresponding eigenvalues, which are proportional to the amount of variance captured by the corresponding PC. To enumerate each principal component, eigenvalues were ranked in descending order of absolute magnitude, with larger magnitudes indicating a greater proportion of the total variance captured by the associated PC. To compute the variance captured per PC for SA-preserving surrogate maps in Figure 3F, we used the following procedure:

1. 50 genes were randomly selected without replacement from a set of brain-specific genes.
2. 40 surrogate gene expression maps were constructed for each selected gene.
3. 1,960 surrogate maps (the number of brain-specific genes) were randomly selected without replacement from the set of 50 × 40 = 2, 000 surrogate maps.
4. PCA was performed on these gene expression maps to determine the ten leading PCs and their variance spectrum.

To construct the variance captured per PC for randomly shuffled surrogate maps, for each replicate, we randomly permuted all 1,960 brain-specific gene expression maps.

### Data visualization

Cortical surface-based visualizations of empirical and surrogate brain maps were generated using the Connectome Workbench software (Glasser et al., 2016b). Left-hemispheric cortical data were illustrated on either flat, spherical, or very inflated cortical surface meshes in the HCP and Conte69 atlases. Cerebellar flat maps were generated using the SUIT Matlab toolbox (http://www.diedrichsenlab.org/imaging/suit.htm) (Diedrichsen and Zotow, 2015) using custom Python scripts adapted from Guell et al. (2018). The black trend line in Fig. 3C was calculated using the Theil-Sen estimator, a nonparametric estimator of linear slope that is insensitive to the underlying distribution and robust to statistical outliers (Sen, 1968).

### Moran spectral randomization

Surrogate maps derived via Moran spectral randomization (MSR) were generated using the singleton procedure implemented in the BrainSpace toolbox (de Wael et al., 2019). Our spatial weight matrix was constructed by inverting the parcellated geodesic distance matrix and setting diagonal elements to 1. The tolerance (i.e., the minimum value for an eigenvalue to be considered non-zero) was set to 10^-6^ as in the online tutorials (http://brainspace.readthedocs.io). The example target brain map was constructed by superimposing normally distributed noise with an exponentially decaying component parametrized by the distance to area MT:

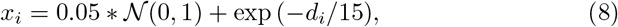

where *x_i_* is the value of the brain map in parcel *i*, and *d_i_* is geodesic distance (in millimeters) of parcel *i* from area MT.

## Results

Physical and mathematical properties of signal-generating processes in the brain induce SA in empirical brain maps (Chumbley and Friston, 2009). To illustrate the statistical impact of SA in brain maps, we consider two MRI-derived structural neuroimaging maps: the T1w/T2w map, which partly reflects intracortical gray-matter myelin content (Glasser and Van Essen, 2011; Glasser et al., 2014), and the cortical thickness map (Fig. 1A). In these two cortical maps, proximal brain regions exhibit more similar values than pairs of spatially distant regions. This property differs starkly with the randomly shuffled (i.e., permuted) brain maps in Figure 1B: randomly shuffling a brain map necessarily destroys its SA structure.

**Figure 1:**
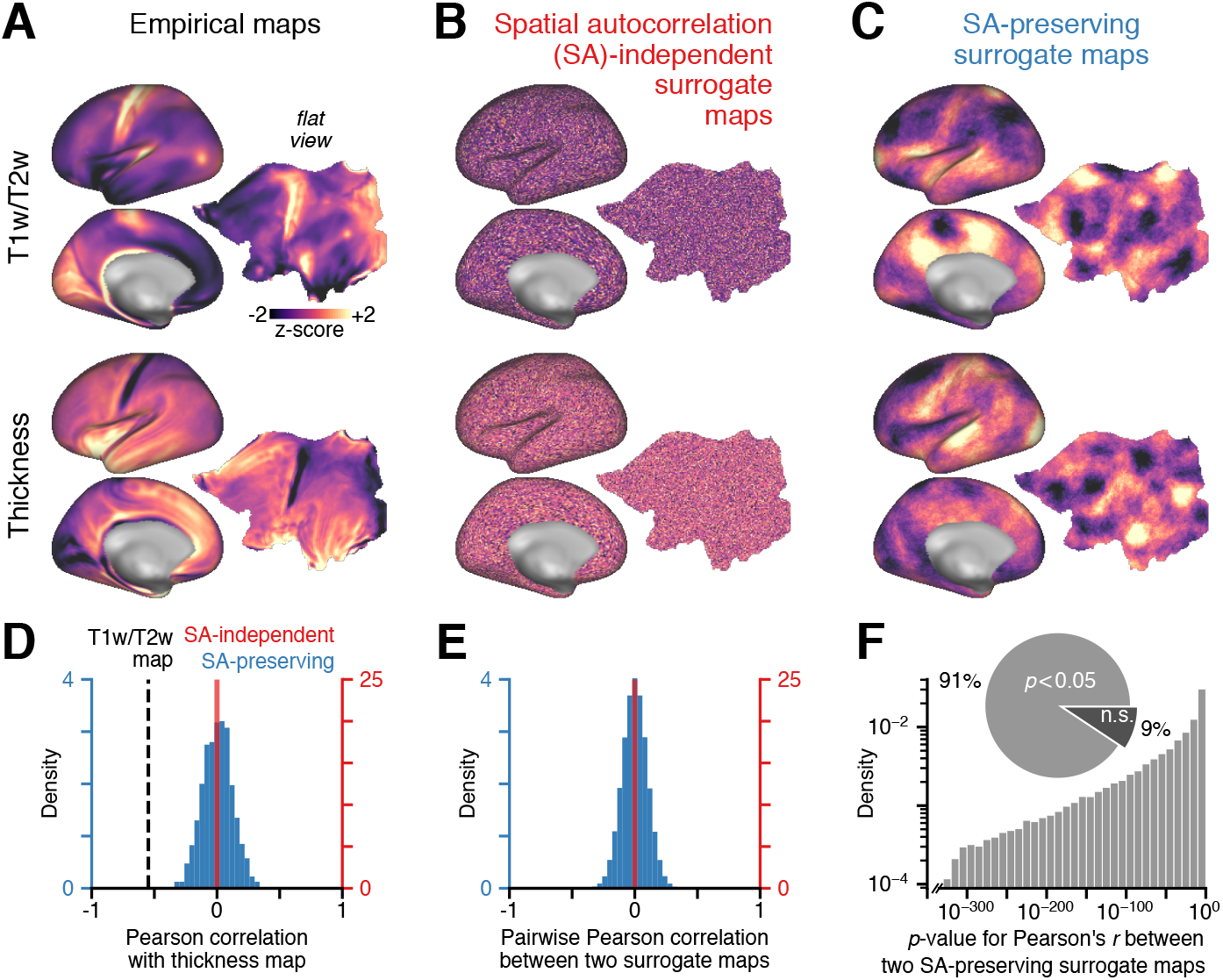
Spatial autocorrelation (SA) in empirical brain maps has a substantial impact on statistical inference. (**A**) The group-averaged (*N* = 339) T1w/T2w (top) and thickness (bottom) maps in the left cortical hemisphere. The maps are spatially autocorrelated: proximal brain regions exhibit more similar values than pairs of spatially distant regions. (**B**) Randomly shuffling the empirical maps — equivalent to assuming samples’ exchangeability, a more relaxed assumption than independence — destroys their autocorrelation structure. (**C**) One example realization of random SA-preserving surrogate maps, derived for each empirical map. (**D**) Null distributions of Pearson correlation coefficients between the empirical cortical thickness map, and 1, 000 randomly shuffled (red) and SA-preserving (blue) surrogate maps derived from the empirical T1w/T2w map. Dashed black line indicates the empirical correlation between the T1w/T2w and cortical thickness maps. (**E**) Distributions of Pearson correlation coefficients between pairs of randomly shuffled (red) and SA-preserving (blue) surrogate maps derived from the empirical T1w/T2w map. (**F**) The distribution of naive *p*-values for the Pearson correlation between pairs of SA-preserving surrogate maps derived from the empirical T1w/T2w map. These naive *p*-values are derived under the assumption that samples are independent and normally distributed, in which case the sampling distribution of Pearson’s *r* is a *t*-distribution with *n* – 2 = 29, 694 degrees of freedom. n.s.: not significant.

Although SA is a prominent and ubiquitous property of brain maps, many conventional parametric statistical tests commonly applied to them assume that data points are independent. This is related to the assumption that data points are exchangeable when performing a non-parametric permutation test. In permutation tests of the significance of a brain map topography, null distributions are constructed by repeatedly shuffling the target brain map, as in Figure 1B, and recomputing the test statistic on these maps. This process preserves the map’s distribution of values while randomizing its topography.

In statistical tests which do not account for the intrinsic SA structure of brain maps, the null hypothesis is that unstructured maps, like those in Figure 1B, are reasonably likely to have produced a comparable or more extreme statistical measure. To increase our confidence in claims regarding the specific spatial topography of a brain map, we propose an alternative null hypothesis, in which null distributions are derived from surrogate brain maps that preserve empirical SA while randomizing topography. By construction, SA-preserving surrogate brain maps would preserve the two-point autocorrelation among brain regions (Fig. 1C).

We found that null distributions for the Pearson correlation coefficient were substantially different when derived from randomly shuffled and SA-preserving surrogate maps. We constructed these distributions by correlating the empirical cortical thickness map with maps in two different sets of surrogates—randomly shuffled and SA-preserving, each derived from the empirical T1w/T2w map (Fig. 1D). SA-preserving surrogate maps produced a null distribution whose variance was more than an order of magnitude greater than the variance of the null distribution produced by randomly shuffled surrogate maps. A similarly large difference in null distribution variance was also found for parcellated brain maps (Supplementary Fig. 2).

This contrast between these approaches is recapitulated in the distributions of pairwise correlations between pairs of surrogate maps (Fig. 1E). As in Figure 1D, the variance of these distributions provides a measure of variation across surrogate maps under the two respective null hypotheses. Increased null distribution variance for the SA-preserving surrogate maps indicates that these maps are, on average, more statistically similar than their randomly shuffled counterparts. Moreover, Pearson correlation *p*-values, which are blind to SA structure, tend to be exceptionally small when computed between pairs of random autocorrelated maps (Fig. 1F): we found that 91% of all pairwise correlations between SA-preserving surrogate maps were statistically significant when assessed using the naive Pearson correlation threshold of *p* < 0.05. This shows that even two randomly generated brain maps are highly likely to be significantly correlated when evaluated using SA-naive statistical measures. Together, these findings demonstrate the substantial impact that SA has on statistical measures derived from large-scale brain maps.

### Constructing spatial autocorrelation-preserving surrogate maps

Figure 2 provides a schematic of our generative modeling method for SA-preserving surrogate brain maps. The target brain map is first permuted, randomizing the map’s topography. Next, SA is reintroduced by smoothing the permuted map with a distance-dependent kernel. Motivated by previous work (Burt et al., 2018; Romero-Garcia et al., 2018; Markov et al., 2011; ArnatkevicIūtė et al., 2019), here we use a smoothing kernel with weights which fall off exponentially with distance. However, we found that null distributions are largely insensitive to the functional form of the kernel (Supplementary Fig. 3). The kernel is truncated at the *k*-th nearest neighbor — differences in this parameter correspond to differences in the characteristic length scale of the autocorrelation which is reintroduced.

**Figure 2:**
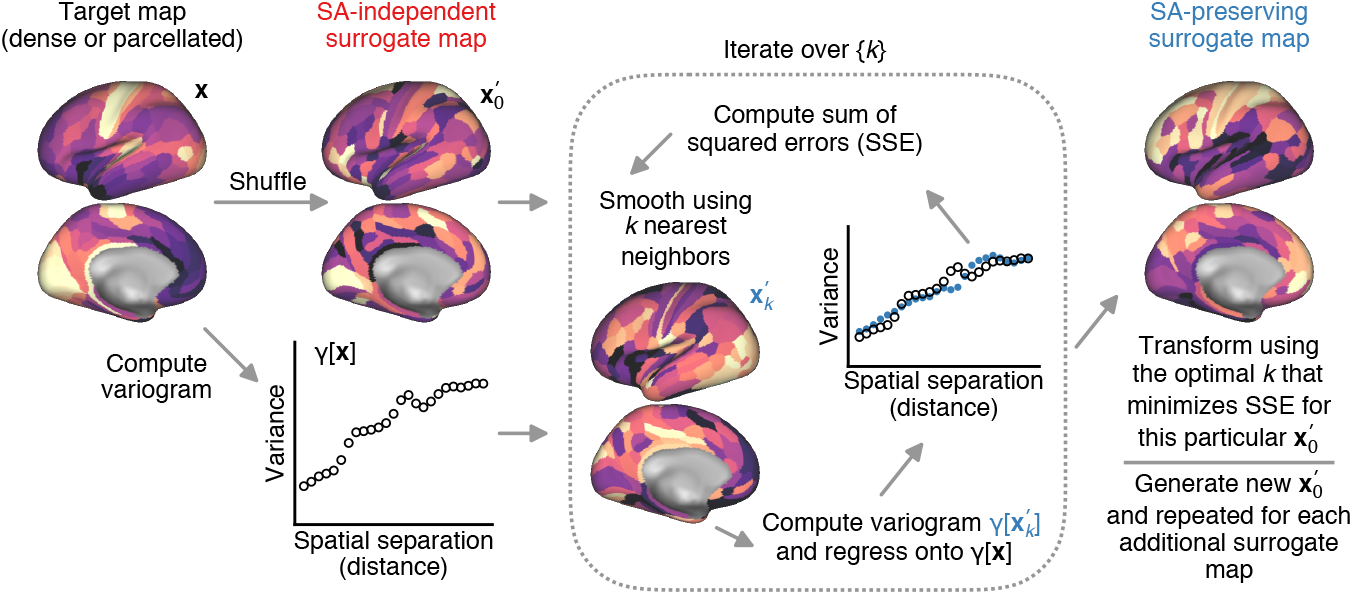
Generating SA-preserving surrogate maps. A dense or parcellated target brain map (x) is first randomly shuffled 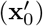, destroying its SA. SA is reintroduced by smoothing the shuffled map with an exponentially decaying, distance-dependent smoothing kernel which includes the *k* nearest neighbors to each region 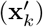. Variograms (*γ*[·]), which we use to operationalize SA, are computed for the target map and the smoothed map. The variogram for the smoothed map is regressed onto the variogram for the target map. The regression coefficients define the transformation of 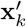 which approximately recovers the autocorrelation in x. The smoothing and regression steps are repeated, each time with a different number of nearest neighbors, *k*, used to perform the spatial smoothing. For each iteration, the sum of squared error (SSE) in the variogram fit is computed; the regression coefficients for the best value of *k* which minimizes SSE are then used to produce a surrogate map whose SA is most closely matched to the target map’s SA.

We operationalize SA in a brain map by computing its variogram. The variogram provides a summary measure of pairwise variation in a map as a function of distance. For instance, any sequence of independent and identically-distributed random variables has no distance dependence in its variation and therefore has a flat variogram. In contrast, a positively autocorrelated spatial map has a variogram with positive slope, because variation among regions in close spatial proximity (at small distances) is less than variation among widely separated regions (at large distances). We construct variograms by computing the brain map’s sample variance within uniformly spaced distance intervals for both the target map and the smoothed map (Equation 2). For surface-based maps, we calculate distance between brain regions using surface-based geodesic distance, while for volumetric maps we use three-dimensional Euclidean distance.

To recover SA in surrogate brain maps, we first perform a linear regression between the smoothed map’s variogram and the target map’s variogram. We then compute the sum of squared error (SSE) in the variogram fit: smaller SSE corresponds to improved recovery of SA structure. To maximize the recovery of SA, the sequence of steps described above — smoothing the permuted map, fitting the variogram, and computing SSE — are repeatedly performed, each time using a different number of nearest neighbors, *k*, used to smooth the permuted map. Finally, the best value of *k*, denoted *k*^*^, which minimizes SSE is used to construct the SA-preserving surrogate map (via Equation 6).

The assumptions of this surrogate generation procedure are that maps are normally distributed and stationary (Viladomat et al., 2014). Convolutions of data and Gaussian smoothing of images, which are standard in neuroimaging data processing pipelines, result in maps which are approximately normal, per central limit theory. Regardless, the value distributions of brain maps and surrogate maps can be invertibly transformed prior to, and following, the use of our method.

### Illustrative applications to empirical neuroimaging data

To demonstrate how SA influences statistical outcomes, we used our method to evaluate statistical significance for three familiar types of brain map analyses: testing for the functional network specificity of a brain map; testing the topographic alignment between two maps; and testing data dimensionality. In each analysis, we compare our findings to the results of standard spatially naive statistical approaches. We perform each test on parcellated brain maps, where neuroanatomically informed parcel borders should mitigate the impact of SA, yielding more conservative estimates for the discrepancies between spatially naive and SA-preserving approaches.

First, we consider the functional network specificity of the MRI-derived cortical T1w/T2w map (Fig. 3A,B). A spatially naive Wilcoxon signed-rank test suggests that mean T1w/T2w map value is significantly higher in sensory networks than in association networks. Like the Wilcoxon signed-rank test, a spatially naive permutation test produces a highly significant *p*-value (*p* < 10^-4^; note that this is a conservative upper bound constrained by the number of permutations). However, functionally specialized networks of brain regions (and regions of interest in general) tend to be spatially contiguous. We therefore computed null distributions of network specificity (i.e., mean map value in association vs. sensory networks) derived from SA-preserving surrogate maps to determine the distribution of results expected by chance. We found that functional network specificity of the cortical T1w/T2w map remains statistically significant, but that the calculated *p*-value is highly attenuated (*p* = 0.01), reflecting the more stringent null hypothesis that the specificity can be driven by SA-constrained maps exhibiting random topography.

**Figure 3:**
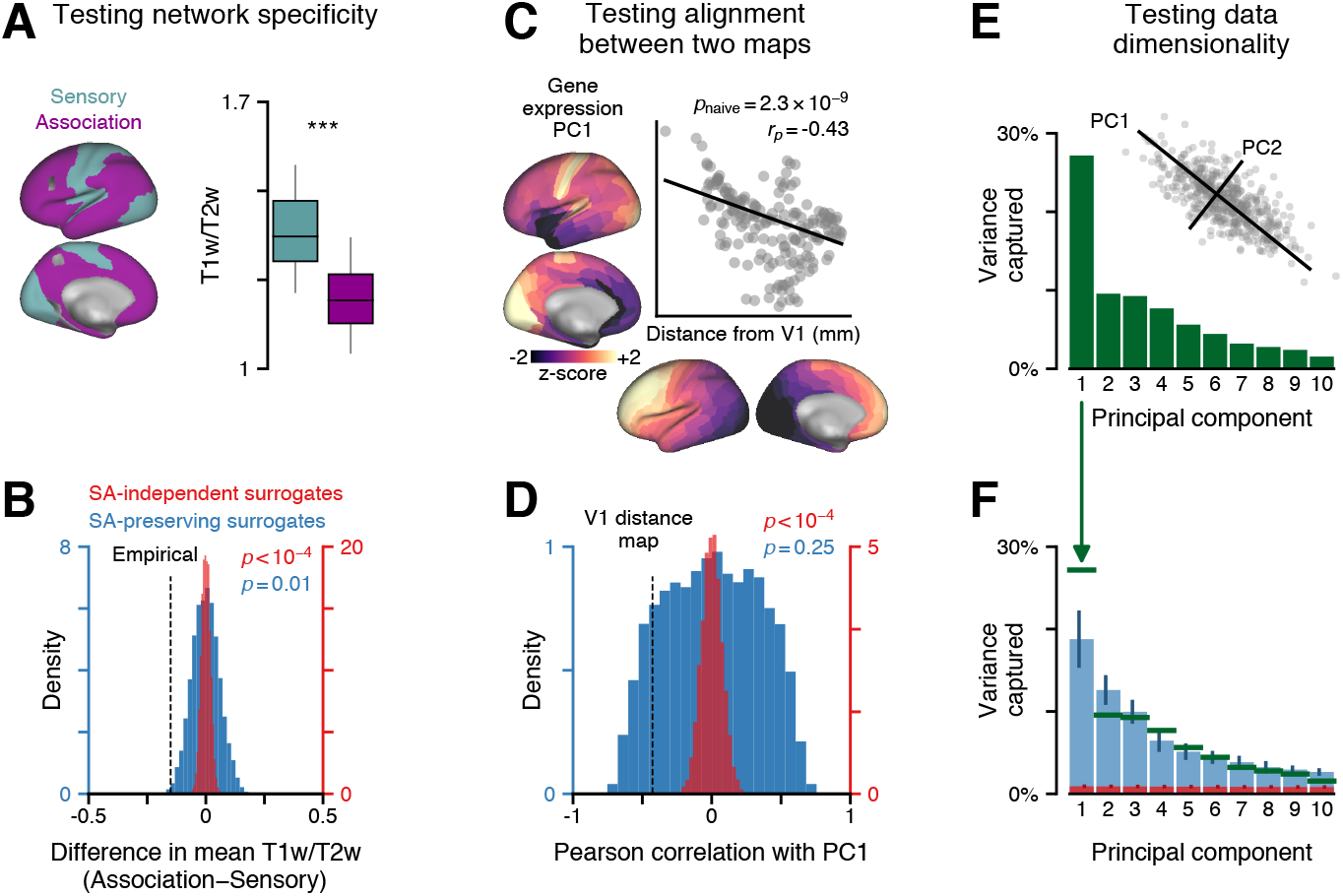
SA-preserving surrogate maps provide a more conservative and meaningful measure of statistical significance in conventional neuroimaging analyses. (**A**) Mean cortical T1w/T2w map value in functionally-defined sensory and association brain networks, computed across 180 parcels in the left cortical hemisphere. Spatially naive statistics suggest that cortical T1w/T2w map value is significantly higher in sensory networks (***; *p* < machine precision; two-sided Wilcoxon signed-rank test). Box plots mark the median and inner quartile ranges across regions within each network, and whiskers indicate the 95% confidence interval. (**B**) Null distributions for the difference in mean T1w/T2w map value between sensory and association networks, derived from 10, 000 randomly shuffled (red) and SA-preserving (blue) surrogate brain maps. Dashed black line indicates the empirically observed difference. (**C**) Spatially naive statistics suggest that the leading principal component (PC1) of cortical gene expression variation, and the map of geodesic distance from visual area V1, are significantly correlated (*r_p_* = — 0.43, *p* = 2.3×10^-9^; Pearson correlation). (**D**) Null distributions of Pearson correlations between gene expression PC1 and 10, 000 randomly shuffled (red) and SA-preserving (blue) surrogate maps, derived from the map of distance from V1. Dashed black line indicates the empirically observed correlation. (**E**) The spectrum of variance captured by the first ten spatial PCs of cortical gene expression variation. PC1 captures a disproportionately large fraction of gene expression variance, indicating that cortical gene expression variation is quasi-one dimensional. (**F**) Null distributions of variance captured per PC, derived by performing PCA on ten replicates of randomly shuffled (red) and SA-preserving (blue) surrogate gene expression maps. Vertical bars indicate standard deviation across replicates. Green horizontal bars indicate the empirical variance spectrum.

Next, we consider the commonly examined problem of assessing correspondence or spatial alignment between two brain maps (Fig. 3C,D). In Burt et al. (2018), we showed that the first principal component (PC1) of brain-specific gene expression variation exhibits a spatial topography that is strikingly similar to the cortical T1w/T2w map — a map which we showed provides a robust noninvasive correlate of cortical hierarchy. Surface-based geodesic distance from primary visual area V1 has also been proposed as proxy measure of cortical hierarchy (Wagstyl et al., 2015) (Fig. 3C). We revisited one of our prior analyses (Burt et al., 2018) and asked whether the map of geodesic distance from area V1, like the T1w/T2w map, is strongly associated with PC1 of gene expression variation. A spatially naive Pearson correlation computed between these two brain maps suggests that their relationship is highly significant (*r_p_* = −0.43; *p* = 2.3 × 10^9^). However, when PC1 is correlated with SA-preserving surrogate maps, each derived from the V1 distance map, the resulting null distribution reveals that this seemingly strong relationship can be explained by SA structure alone (*p* = 0.25; Fig. 3D).

Finally, we examine how SA in brain maps influences principal component analysis (PCA), a common linear decomposition and dimensionality reduction technique. PCA identifies an orthogonal decomposition of data into dimensions along which the data principally vary. When principal components (PCs) are rank-ordered according to the amount of variance they capture (i.e., such that PC1 captures more variance than PC2), the shape of the distribution (i.e., the variance spectrum) provides information about the data’s dimensionality. In Burt et al. (2018), we found that PC1 of brain-specific gene expression variation captures an appreciable fraction of gene expression variance (Fig. 3E), indicating that gene expression primarily varies along the spatial mode defined by PC1’s topography.

To determine whether or not the low dimensionality of gene expression variation can be explained simply by SA in the transcriptional data, we performed PCA on ten replicate sets, each comprising 1,960 SA-preserving surrogate gene expression maps (Fig. 3F). We found that the variance captured by empirical PC1 greatly exceeds the expected variance captured by chance for spatially autocorrelated surrogate maps, while subsequent empirical PCs exhibit a pattern of captured variance which closely matches the null variance spectrum. In contrast, PCA performed on replicate sets of randomly shuffled gene expression maps yields a much flatter null variance spectrum. These discrepancies are due to differences in how variance is redistributed: random shuffling tends to redistribute variance uniformly across the brain, whereas SA-preservation retains the low-dimensional structure of variance found in the empirical data.

Collectively, these findings underscore the need for plausible null hypotheses, and associated principled null models, to properly evaluate statistical outcomes when performing tests on brain maps with large-scale spatial gradients. In each analysis, incorporating SA directly into the null hypothesis had a substantial impact on inference.

### Spatial autocorrelation modulates null distribution variance

To further characterize the impact of SA on statistical measures, we investigated the relationships between SA structure and null distribution variance. We first considered the simplified mathematical setting of Gaussian random fields (GRFs), which are random fields with a multivariate normal probability distribution. The SA structure of a GRF can be parametrically controlled by changing the slope of the field’s power spectral density (Appendix B). We define our field’s power spectra to have the functional form *P*(*k*) = |*k*|^-α^, where *α* > 0 and *k* is a spatial frequency (not to be confused with the number of nearest neighboring regions). Intuitively, as *α* increases, spectral power *P* becomes increasingly concentrated at low spatial frequencies, yielding increasingly autocorrelated fields.

Three realizations of GRFs with varying SA structure are illustrated in Figure 4A. Differences in the fields’ SA are reflected in the shapes of their variograms: fields with greater SA are less variable across greater distances (Fig. 4B). To relate SA to null distribution variance, for each GRF realization we constructed *N* = 1, 000 SA-preserving surrogate fields, and computed distributions of pairwise Pearson correlation coefficients between fields at each SA level (Fig. 4C). We found that null distribution width (i.e., variance) increased as a function of SA (Fig. 4D), suggesting that SA tends to reduce the number of effective degrees of freedom (i.e., the number of ways in which the fields can vary). To understand this phenomenon intuitively, consider the limiting case in which a one-dimensional random field is perfectly autocorrelated. In this limit, the system reduces to a single degree of freedom, i.e., lines with variable slopes. Thus, the distribution of pairwise Pearson correlations between these lines would be a bimodally peaked “distribution” with equal probabilities of obtaining *r_p_* = ±1, depending on whether the slopes have equal or opposite signs—in other words, a distribution with maximal variance.

**Figure 4:**
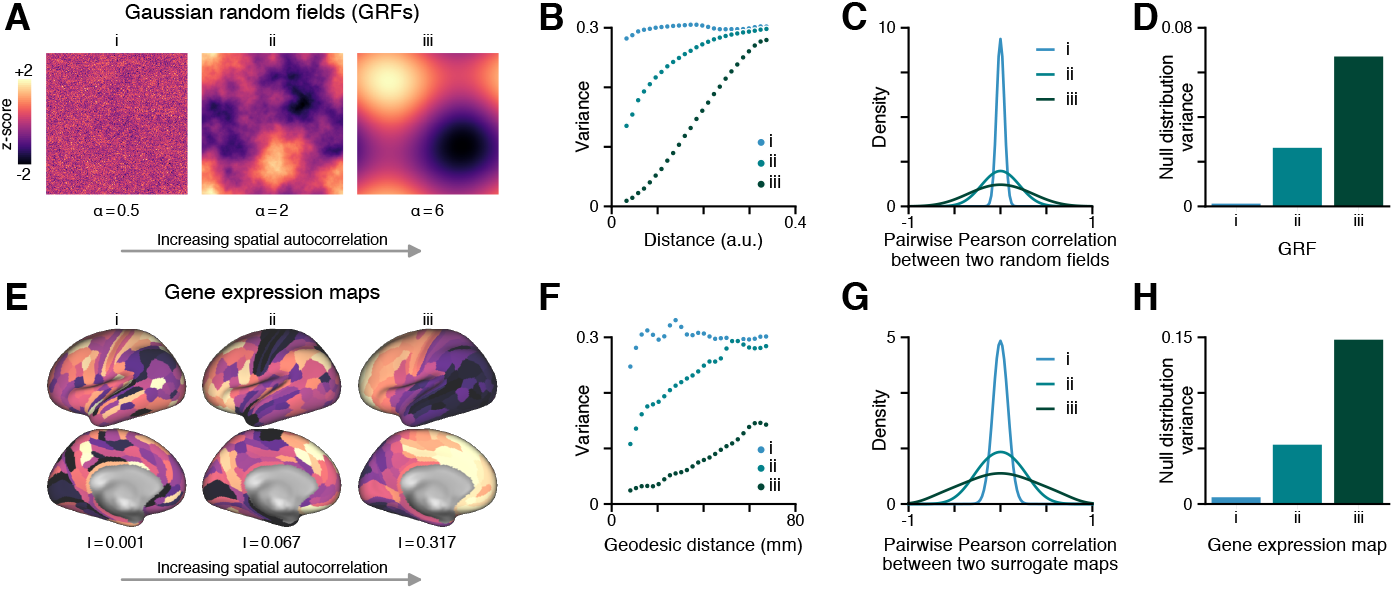
SA drives an increase in null distribution width. (**A**) Gaussian random fields (GRFs) with increasing SA, simulated on a uniformly spaced lattice of 100^2^ points. SA is varied by changing the slope of the fields’ power spectral density, *P* (*k*) = |*k*|^-*α*^. (**B**) Variograms for the three GRFs in **A**. SA reduces variability between spatially proximal points. (**C**) Distributions of pairwise Pearson correlations between 1, 000 random realizations of each field in **A**. (**D**) Variance of each distribution in **C**. SA constrains the fields’ variability such that their pairwise correlations tend to be larger in magnitude. (**E**) Gene expression variation across 180 parcels in the left cortical hemisphere. Moran’s *I* statistic, a measure of SA, was used to select three genes whose cortical expression maps exhibit low (i), moderate (ii), and high (iii) SA structure. (**F**) Variograms for the three gene expression maps in **E**. (**G**) Distributions of pairwise Pearson correlations between 1, 000 SA-preserving surrogate maps, computed for each map in **E**. (**H**) Variance of each distribution in **G**. Histograms in panels **C** and **F** were smoothed using Gaussian kernel density estimation. Labels i, ii, and iii in panels **E-H** correspond to genes CCDC18-AS1, SERPINI1, and PRRX1, respectively.

Equipped with these intuitions, we asked whether the same relationship appears in analyses of empirical brain maps with different SA. Because SA in gene expression maps is highly variable (Gryglewski et al., 2018), we repeated the analyses described above using parcellated gene expression maps derived from microarray data in the Allen Human Brain Atlas (Burt et al., 2018; Hawrylycz et al., 2012, 2015). To quantify SA in gene expression maps, we computed Moran’s *I* statistic (Moran, 1950). Moran’s I provides a measure of global autocorrelation in spatial data and ranges between −1 and 1, with positive values indicating the presence of positive spatial autocorrelation (i.e., indicating that proximal regions tend to be positively correlated). We then selected three genes, two at the extremes and one at the center of the *I*-distribution (range 0.001–0.317), respectively characterized by low, moderate, and high SA (Fig. 4E). Variograms for these three gene expression maps followed the same trend observed for GRFs (Fig. 4F), and null distribution variance was greater for genes with larger *I* values (with higher SA) (Fig. 4G-H). These results indicate that SA modulates the variance of distributions for both GRFs as well as empirical brain maps. This relationship between SA and null distribution variance reveals the origin of the large discrepancies between *p*-values derived from spatially informed and spatially naive approaches.

### Sampling density and autocorrelation increase the likelihood of type I errors

If a measured brain feature is not spatially autocorrelated, or if it is weakly autocorrelated but only sparsely sampled, then samples should be approximately independent. In this scenario, we expect conventional tests to agree with spatial statistical approaches. In contrast, if a brain feature is strongly autocorrelated, then we expect that as the sampling density is increased, the assumption of statistical independence will be increasingly violated and conventional tests should diverge from spatial approaches. To test this, we investigated whether independently increasing sampling density or SA amplified the discrepancy between spatially informed and spatially naive approaches.

We first demonstrated that our surrogate map generating process is robust across a broad range of spatial scales. Using the HCP’s multi-modal parcellation (Glasser et al., 2016a), dense and parcellated data differ in spatial resolution by two orders of magnitude. Figure 5A-B illustrates that simulated surrogate maps derived from dense and parcellated brain data each reliably recover empirical SA.

**Figure 5:**
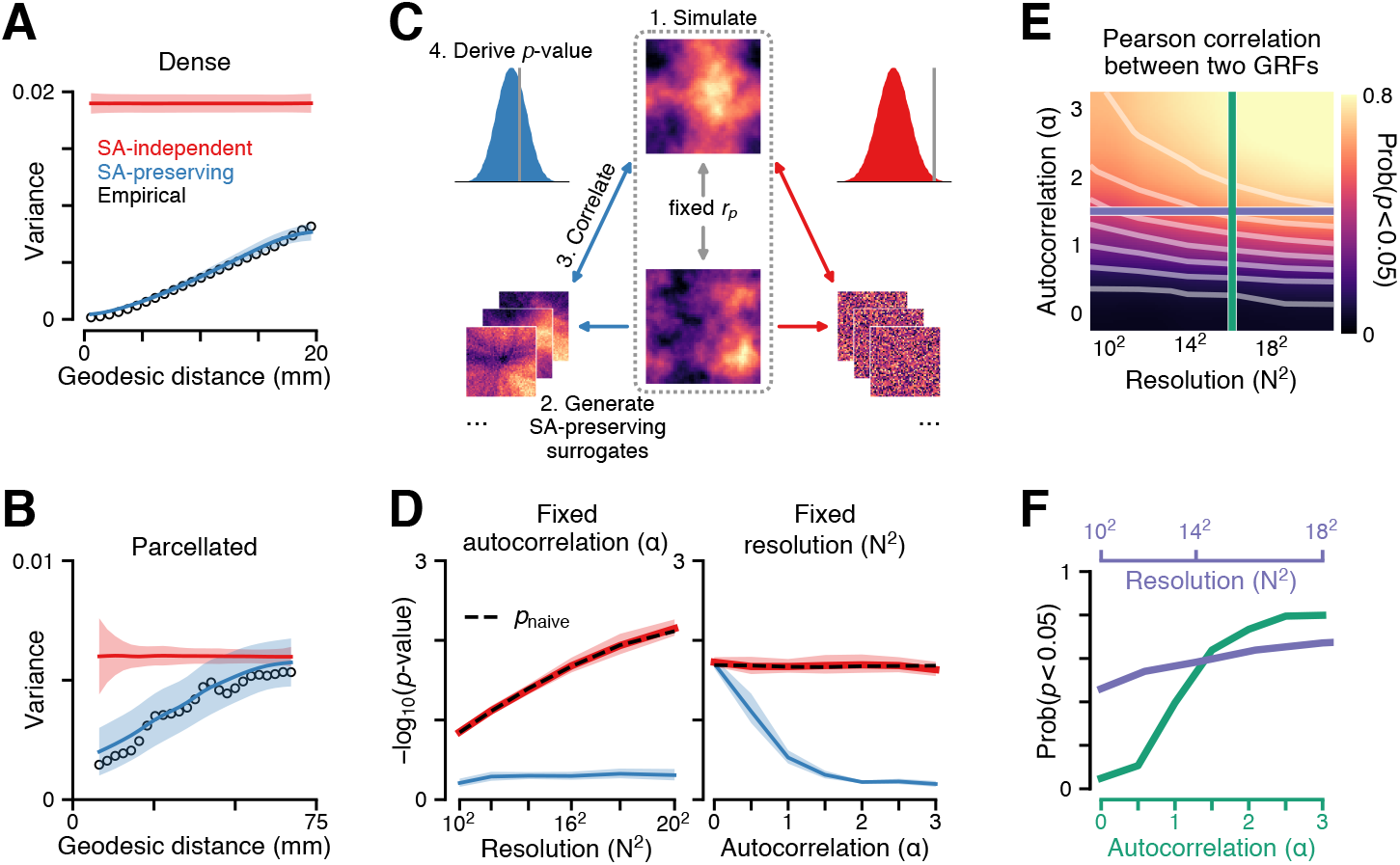
Conventional *p*-values derived from spatially naive tests are highly sensitive to SA and spatial resolution. (**A-B**) Variograms for 1, 000 SA-preserving surrogate maps, derived from the dense (**A**) and parcellated (**B**) group-averaged cortical T1w/T2w map. Open black circles indicate the empirical map’s variogram. Colored lines and shaded regions indicate mean and standard deviation across surrogates. (**C-D**) Pairs of GRFs are simulated until obtaining a pair with a Pearson correlation of |*r_p_*| = 0.15±0.005. Null distributions are computed between one of the fields and 1, 000 randomly shuffled (red) and SA-preserving (blue) surrogate fields, derived from the second field in the pair. Statistical significance is assessed three ways: a parametric *p*-value is derived from a Student’s *t*-distribution with *N*^2^ −2 degrees of freedom (black), and non-parametric *p*-values are derived from the two null distributions. Colored lines and shaded regions indicate mean and standard deviation across 100 replicates. Left: SA fixed at *α* = 2. Right: Spatial resolution fixed at *N*^2^ = 256. Larger values indicate increased statistical significance. (**E-F**) The probability of obtaining a significant (*p* < 0.05) Pearson correlation between two GRF realizations as a function of SA and resolution. Each data point indicates the mean number of significant comparisons across 5, 000 trials. White contours from bottom to top correspond to probabilities 0.1, 0.2, …, 0.7. Purple and green slices through **E** correspond to lines in **F**.

We then performed our surrogate map analyses on simulated GRFs, for which the SA and grid resolution (a measure of sampling density) can be easily and independently varied (Fig. 5C). First, we simulated a random pair of GRFs with a fixed pairwise Pearson correlation. For one of the two GRFs, we generated both SA-preserving and randomly shuffled surrogate fields. These surrogate fields were then correlated with the second GRF, yielding two null distributions of expected Pearson correlations. We assessed the statistical significance of the association between the two fields in three ways: using a spatially naive parametric *p*-value derived from a Student’s *t*-distribution, which assumes that samples are independent and normally distributed; and by deriving nonparametric *p*-values from the two null distributions.

We determined how these three *p*-values were influenced when independently varying the grid resolution (at fixed SA) and varying the SA (at fixed resolution) (Fig. 5D). Consistent with our hypothesis, we found that the discrepancy between spatially naive and spatially informed approaches grows in magnitude as a function of both resolution and SA: at fixed SA, greater spatial resolution leads to more significant *p*-values for the spatially naive statistical tests, whereas *p*-values derived from SA-preserving surrogate maps remain stable. In contrast, at fixed spatial resolution, increased SA leads to less significant *p*-values for spatially informed tests, whereas it has no effect on the result of spatially naive approaches (because the correlation between the simulated GRF pairs was fixed).

To characterize how these two properties interact, we computed the probability of obtaining a statistically significant (*p* < 0.05) Pearson correlation between two GRFs while jointly varying SA and resolution (Fig. 5E,F). We found that the probability of obtaining a significant result scales with sampling density, and that the strength of this scaling is modulated by the strength of autocorrelation—evidence for an interaction between these two properties. These findings collectively indicate that greater SA and sampling density of autocorrelated processes both independently and jointly increase the likelihood of obtaining type I errors when using conventional tests.

### Constructing subcortical surrogate maps with spatial autocorrelation

To the best of our knowledge, there is currently no established method for generating spatially matched surrogate data for subcortical, or other volumetric, brain maps. To further demonstrate the flexibility of our approach, we generated SA-preserving surrogate maps for a recently identified functional gradient map in human cerebellum (Guell et al., 2018) (Fig. 6A,D). We follow the same procedure to generate our cerebellar surrogates, with the only difference being the distance metric: whereas for cortical surface surrogate maps, we used surface-based geodesic distance between regions, here for our subcortical volumetric surrogate maps we used three-dimensional Euclidean distance. By construction, our generative model produces surrogate cerebellar maps which preserve the empirical SA in the functional gradient map (Fig. 6C,F,I). As in cortex, SA-preserving cerebellar surrogate maps produce null distributions with considerably more variance than null distributions constructed from randomly shuffled maps (Fig. 6G,H).

**Figure 6:**
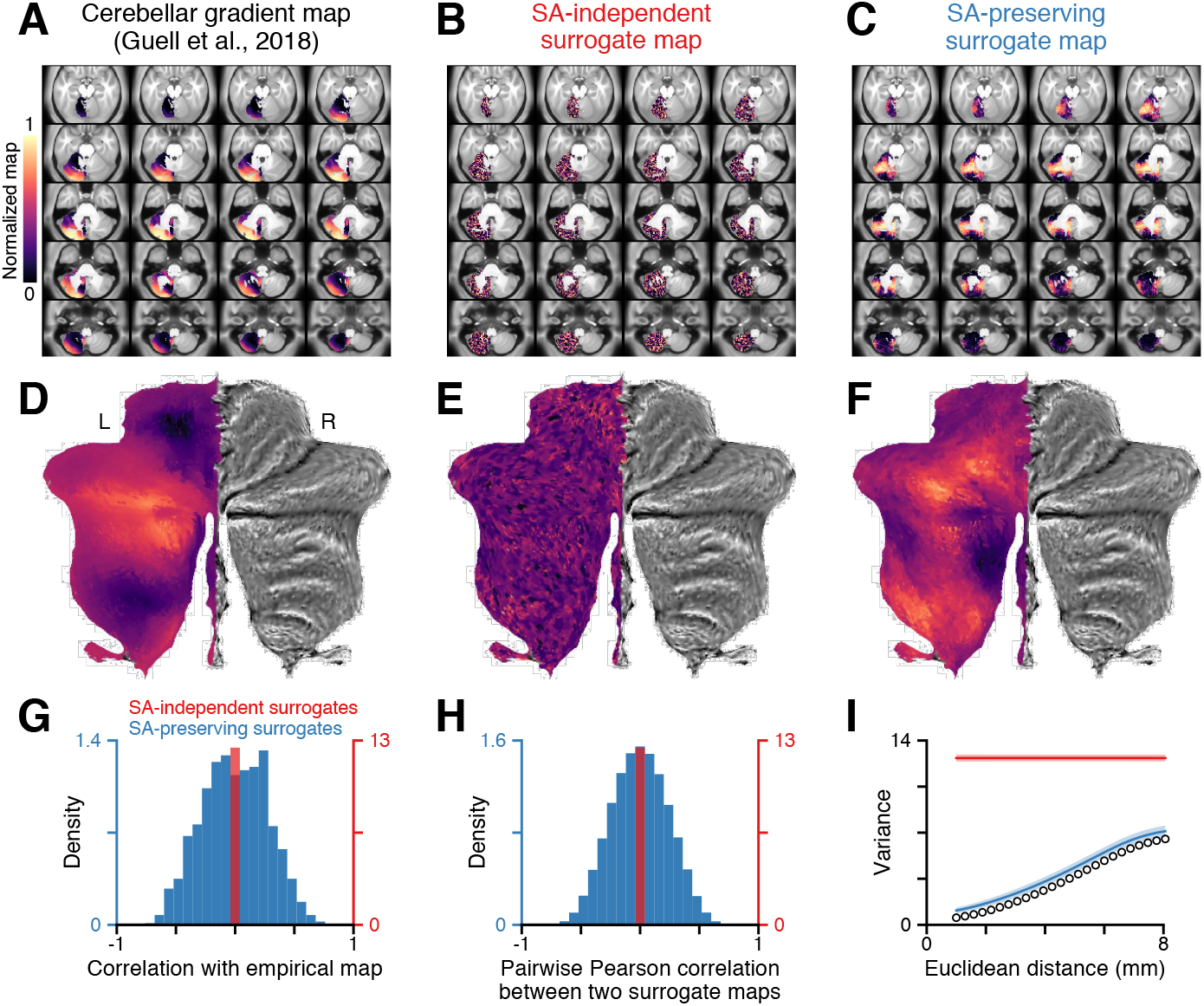
Generating SA-preserving surrogates for a subcortical map. (**A**) Functional cerebellar gradient 1 map, derived from diffusion map embedding of resting-state functional connectivity data by Guell et al. (2018), shown for the left hemisphere. (**B**) Randomly shuffling the map destroys its SA. (**C**) SApreserving volumetric surrogate maps of the cerebellum preserve the SA present in the empirical map. (**D-F**) Flat projections of the volumetric cerebellar maps in **A-C**, constructed using the SUIT toolbox in Matlab. (**G**) Null distributions of Pearson correlation coefficients between the empirical map and 1, 000 randomly shuffled (red) and SA-preserving (blue) surrogate maps, derived from the functional gradient map. (**H**) Distributions of Pearson correlation coefficients between pairs of randomly shuffled (red) and SA-preserving (blue) surrogate maps, derived from the functional gradient map. (**I**) Variograms for the empirical map (open black circles) and 1, 000 randomly shuffled (red) and SApreserving (blue) surrogate cerebellar maps. Colored lines and shaded regions indicate mean and standard deviation across surrogates.

### Spatial autocorrelation drives enrichment in gene ontology analyses

Recent advances in high-throughput transcriptomics have made it possible to perform gene expression profiling throughout the brain and across the genome. Advances in bioinformatics have also produced gene ontology (GO) databases, which provide a corpus of functional annotations (or GO categories) that relate genes to specific biological functions and molecular pathways. Together, transcriptomic profiling and GO databases establish putative mappings between large-scale gene expression topography and biological function.

A growing number of recent studies have leveraged these technologies to infer biological functions associated with large-scale brain maps using GO enrichment analyses (Vértes et al., 2016; Whitaker et al., 2016; Romero-Garcia et al., 2018; Morgan et al., 2019). These map-GO analyses often follow a similar logic (Fig. 7A). First, a brain map of interest is compared to maps of gene expression using uni-or multi-variate approaches. Each gene is ranked according to the strength of the association between its expression profile and the brain map’s spatial topography. A GO enrichment analysis is then performed on the topranking genes: when a significant number of top-ranking genes have a particular annotation, relative to the number of annotations present in a reference set (e.g., the entire genome), then the top-ranking genes are said to be enriched for that annotation. Because the top-ranking genes are strongly associated with the brain map of interest, the enriched annotations are used to infer biological functions associated with that particular brain map’s topography.

**Figure 7:**
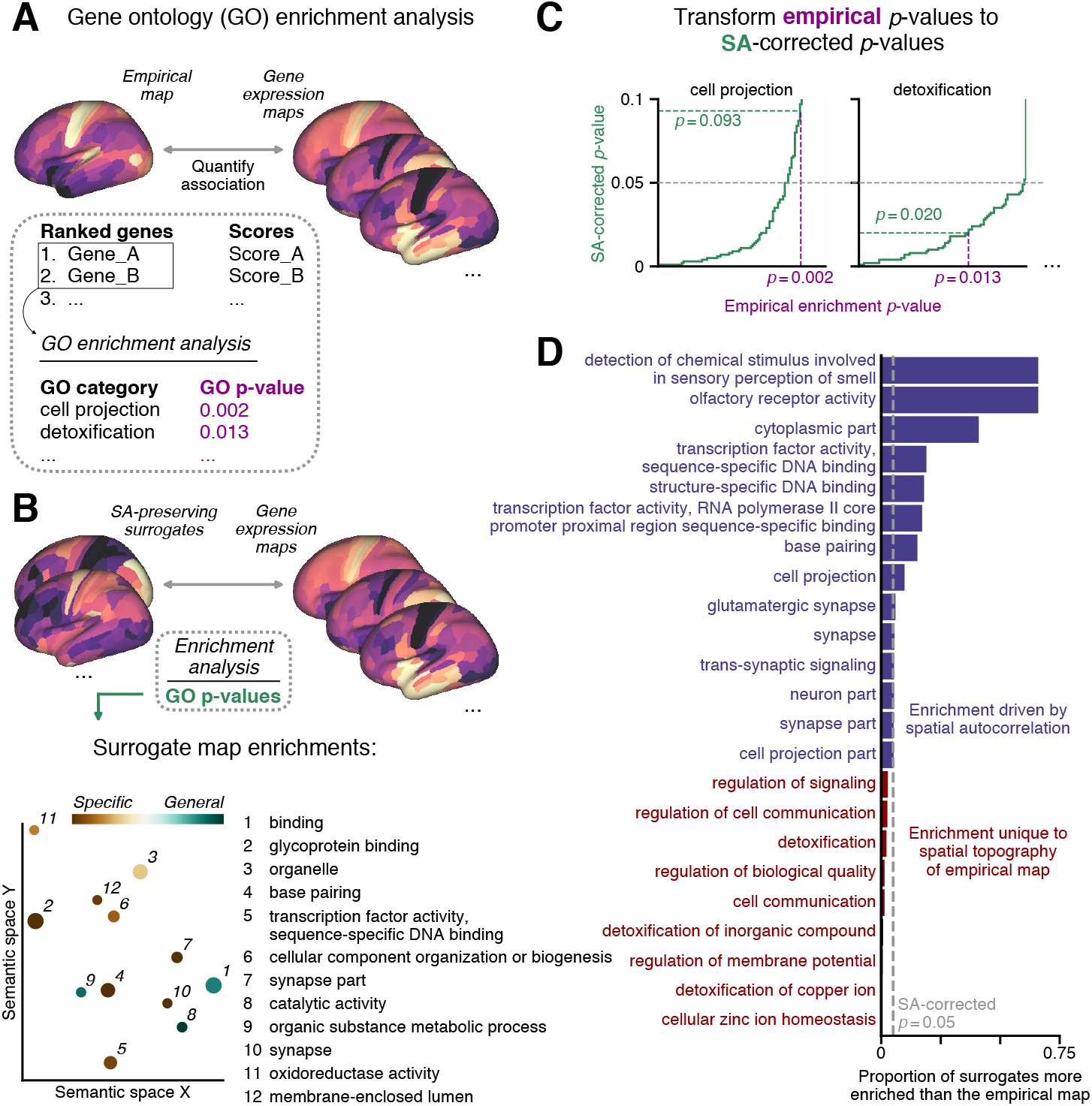
Gene ontology (GO) enrichment analysis of SA-preserving surrogate maps. (**A**) In standard GO enrichment analyses, topographical relationships between a brain map and gene expression maps are first computed using uni-or multi-variate regression techniques, e.g., Pearson correlation or partial least squares regression. Genes are ranked according to their statistical association with the brain map. Significant enrichments — annotations (GO categories) which occur more frequently than expected by chance — for a set of top-ranking genes are then used to infer the biological functions associated with the brain map. (**B**) GO enrichment analysis applied to 1, 000 SA-preserving surrogate maps derived from the cortical T1w/T2w map. Points are scaled in proportion to the fraction of surrogate maps which were significantly enriched (range 13-45%). Points are colored according to the frequency with which each annotation appears across all genes in the genome: specific (general) indicates low (high) frequency (range 0-66%). X- and Y-coordinates derive from multidimensional scaling, such that nearby points are semantically similar, using the web-based tool REViGO. (**C**) SA-preserving null distributions of expected enrichments provide a mapping from empirical enrichment *p*-values (magenta) to surrogate map-corrected *p*-values (green). The point at which the empirical *p*-value (dashed magenta) intersects the cumulative distribution function of surrogate map-derived *p*-values (solid green) indicates the fraction of surrogate maps which were more significantly enriched than the empirical map (dashed green). (**D**) Surrogate map-corrected *p*-values for empirically enriched GO categories. Significant (*p* < 0.05) surrogate map-corrected *p*-values indicate that the empirical map’s enrichment is primarily driven by its specific spatial topography, rather than its statistical properties (SA). Blue (red) bars indicate categories for which more (fewer) than 5% of surrogate maps were more significantly enriched than the empirical T1w/T2w map.

In effect, the null hypothesis in these types of map-GO analyses is that the number of annotations in the top-ranking gene set is expected by chance. In other words, the null hypothesis is that a randomly selected set of genes is expected to contain the number of annotations which were observed empirically. However, both brain maps and gene expression profiles are spatially autocorrelated, and gene expression profiles strongly covary across human cortex (Burt et al., 2018). As a result, not all genes are equally likely to be selected in a GO analysis, regardless of any specific alignment of map topographies. To demonstrate this effect, we performed a GO enrichment analysis on SA-preserving surrogate maps, derived from the cortical T1w/T2w map (Fig. 7B). We found that spatially autocorrelated surrogate brain maps were significantly enriched for many functional annotations (i.e., the top-correlating genes for those surrogate brain maps are significantly enriched). Therefore, under the null hypothesis of conventional map-GO analyses, brain maps which are SA-constrained — yet have random topographies — yield statistically significant GO enrichment.

The above analysis shows that GO enrichment of brain maps is spuriously driven by SA rather than topography. For a brain map of interest, the resulting enriched GO categories may not be a property of that map’s specific topography, but merely its SA. How then can map-GO enrichments be interpreted, for a brain map of interest? We propose a statistical framework which utilizes generative null modeling to test a more stringent and meaningful null hypothesis: that the observed number of annotations is driven by SA in the empirical map, and is therefore not a special property of that brain map’s topography.

Our procedure to test this more meaningful null hypothesis proceeds as follows (Fig. 7B-D). First, SA-preserving surrogate maps are derived from the brain map of interest. For each surrogate map, the top-ranking gene set is computed and then fed into a GO enrichment analysis. After repeating this procedure on all surrogate maps, the result is a null distribution of expected *p*-values for each functional annotation. These distributions establish a mapping from enrichment *p*-values — i.e., the output of the conventional approach — to SA-corrected *p*-values under the new null hypothesis (Fig. 7C). Specifically, SA-corrected *p*-values are derived for each annotation by computing the fraction of surrogate maps which were more significantly enriched than the empirical map. Annotations for which the empirical map is significantly enriched, with respect to its surrogates, correspond to biological functions which are uniquely associated with the empirical map (Fig. 7D). Rejecting this alternative null hypothesis provides much stronger evidence that a brain map has special properties unique to its spatial topography.

To examine the impact of testing this null hypothesis, we applied the procedure to a map-GO analysis for the cortical T1w/T2w map (Fig. 7D). Out of 23 enriched GO categories defined by the conventional enrichment *p* < 0.05, we found that only 9 categories also reached significance with the more stringent SA-corrected *p* < 0.05. That is, enrichment for 14 of the conventionally-selected 23 GO categories could be explained merely by SA of the T1w/T2w map rather than its topography. These findings suggest exercising caution when interpreting map-GO analyses, and demonstrate a procedure to correct for the substantial impact of SA. Furthermore, this generative null modeling framework can be flexibly adapted to other complex statistical analyses of brain maps.

### BrainSMASH: A Python-based platform for simulating surrogate brain maps

We have developed an open-access Python-based computational platform for generating SA-preserving surrogate maps for any brain map of interest (Supplementary Fig. 4). BrainSMASH (Brain Surrogate Maps with Autocorrelated Spatial Heterogeneity) requires only a brain map of interest and a matrix of pairwise distances between elements of the brain map. How these inputs are derived is left to user discretion, though additional support has been provided for investigators working with HCP-compliant surface-based neuroimaging files. In particular, BrainSMASH includes routines to generate two-dimensional Euclidean and geodesic distance matrices from surface geometry (GIFTI) files, and subcortical Euclidean distance matrices from CIFTI-format files. All key parameters described in the methods default to the values used in this study but are easily reconfigurable through the API. Full details are described in the package documentation (https://brainsmash.readthedocs.io).

## Discussion

Here we have presented a method adapted from geostatistics for generative null modeling of surrogate brain maps with SA matched to the SA of a target brain map (Viladomat et al., 2014). We have validated our method and demonstrated its flexibility by showing that simulated surrogate maps recover SA of empirical surface-based and volumetric maps, at parcellated and dense resolutions. Our generative approach makes it possible to formulate and test a specific null hypothesis which accounts for SA, a characteristic and ubiquitous property of brain maps. We have released an open-access Python-based implementation of our method with additional neuroimaging-specific functionality, BrainSMASH.

Studies of large-scale spatial gradients often seek to discover meaningful properties related to the specific topography of a brain map. To do so with confidence, we must distinguish real and meaningful properties from those which can be spuriously driven by general statistical properties of our data, such as SA. In other words, we require methods to determine the likelihood of our observation occurring by chance under a plausible null hypothesis which incorporates general constraints on the space of possible alternatives. In practice, often the choice of null hypothesis is not obvious; rather, it is implicit in the applied statistical test. However, many conventional tests such as the Pearson correlation do not control for SA, which is a prominent feature of brain maps. Incredibly small *p*-values produced by these spatially naive methods should be interpreted not as evidence of significance, but merely as an indication of how poorly the null hypotheses can explain the observations. Spatially naive methods allow one to reject the possibility that unstructured noise, which forms neurobiologically implausible maps (Fig. 1B), can explain the observations. In contrast, SA-preserving methods allow one to reject the possibility that a map with a random topography but comparable SA (Fig. 1C) can explain the observations.

Spatial dependence is an important property of brains: local features and inter-regional associations are influenced by the spatial arrangement of brain regions. For instance, the non-independence of signals measured in proximal brain regions impacts expectations for the spatial extent of task activation peaks in neuroimaging (Friston et al., 1994). In network neuroscience, distance-dependent wiring rules have been incorporated into generative null network models to establish expectations for graph-theoretic measures (Song et al., 2014; Betzel et al., 2016). Here we have incorporated SA into our null hypothesis in an adaptive (i.e., target brain-map specific) manner due to its prominence and ubiquity in brain maps (Burt et al., 2018; Markov et al., 2011; ArnatkevicIūtė et al., 2019; Romero-Garcia et al., 2018), its variation across brain maps (Burt et al., 2018; Gryglewski et al., 2018), and because of its profound impact on statistical measures of interest.

Generative null modeling facilitates hypothesis testing for arbitrarily complex statistical measures. This flexibility considerably broadens their scope of applicability relative to conventional methods for computing *p*-values directly. Generating surrogate data is particularly useful when the sampling distribution of a statistic lacks a closed-form expression. For instance, Demirtaş et al. (2019) used a hierarchical cortical gradient to parametrize a dynamical brain model that simulates functional connectivity, and tested the impact of this gradient relative to SA-preserving surrogates. Surrogate data instantiate an explicitly formulated null hypothesis, and thereby reproduce the expected distribution of a statistic under that hypothesis (Fornito et al., 2016). Generative null models thus have widespread utility as tools for power calculations and statistical inference.

Recent studies have proposed alternative methods to account for SA in statistical analyses of brain maps. The spin test was recently developed to test the anatomical alignment between two cortical surface maps (Alexander-Bloch et al., 2018). The spin test, however, cannot be used to test alignment between volumetric maps or maps which span only a small subset of cortex. Variably sized and irregularly spaced parcels in cortical parcellations also make the spin test impractical for comparisons between parcellated brain maps. Furthermore, the spin test is not suitable for generating SA-preserving surrogate maps (Supplementary Fig. 5). Spatial autoregressive modeling (Burt et al., 2018) and Moran spectral randomization (de Wael et al., 2019) can be used to generate spatially autocorrelated surrogate brain maps. However, these models require the user to choose a specific functional form of the spatial dependence among regions, and may be highly sensitive to this choice (Dubin, 1998). Furthermore, Moran spectral randomization may produce surrogate maps which are strongly correlated with the target brain map, and are therefore not suitable surrogate data for constructing null distributions (Supplementary Fig. 6). In contrast, by construction our method generates surrogate maps which exhibit random topographies.

Generative null modeling is a powerful and flexible approach for evaluating statistical measures against an explicitly defined null hypothesis. The present study, which presents a generative model of brain maps with constrained SA structure, extends our ability to control for a prominent and ubiquitous feature of neuroimaging data. Future work can build on this approach to incorporate additional constraints within a generative modeling framework, thereby expanding the scope of scientific inquiry in the study of large-scale brain organization.

## Acknowledgements

This research was supported by NIH grants R01MH112746 (J.D.M.) and R01MH-108590 (A.A.), a SFARI Pilot Award (J.D.M., A.A.), and BlackThorn Therapeutics (J.D.M., A.A.). The funding sources had no involvement in study design; nor in the collection, analysis, and interpretation of data; nor in the writing of the manuscript; nor in the decision to submit the manuscript for publication.

## Competing Interests

M.H. and M.S. declare no competing interests. J.B.B. consults for Blackthorn Therapeutics. A.A. and J.D.M. consult for and hold equity with Blackthorn Therapeutics.

## Author Contributions

All authors helped design the research. J.B.B. conducted the analyses. J.B.B., M.H., and M.S. wrote the software package. J.D.M. supervised the project. J.B.B. and J.D.M. wrote the manuscript and prepared the figures. All authors contributed to editing the manuscript.

## Author Information

Correspondence and requests for materials should be addressed to J.D.M.

## Code and data availability

A Python-based implementation of the surrogate map generating algorithm used to conduct analyses in this study may be downloaded as an open-access software package, BrainSMASH: https://github.com/murraylab/brainsmash. All results derive from data that are publicly available from sources described above.

## Appendix A. Algorithms for generating surrogate maps

To generate parcellated and dense SA-preserving surrogate maps, we use the following two algorithms:

### Algorithm 1 Generating a parcellated SA-preserving surrogate map

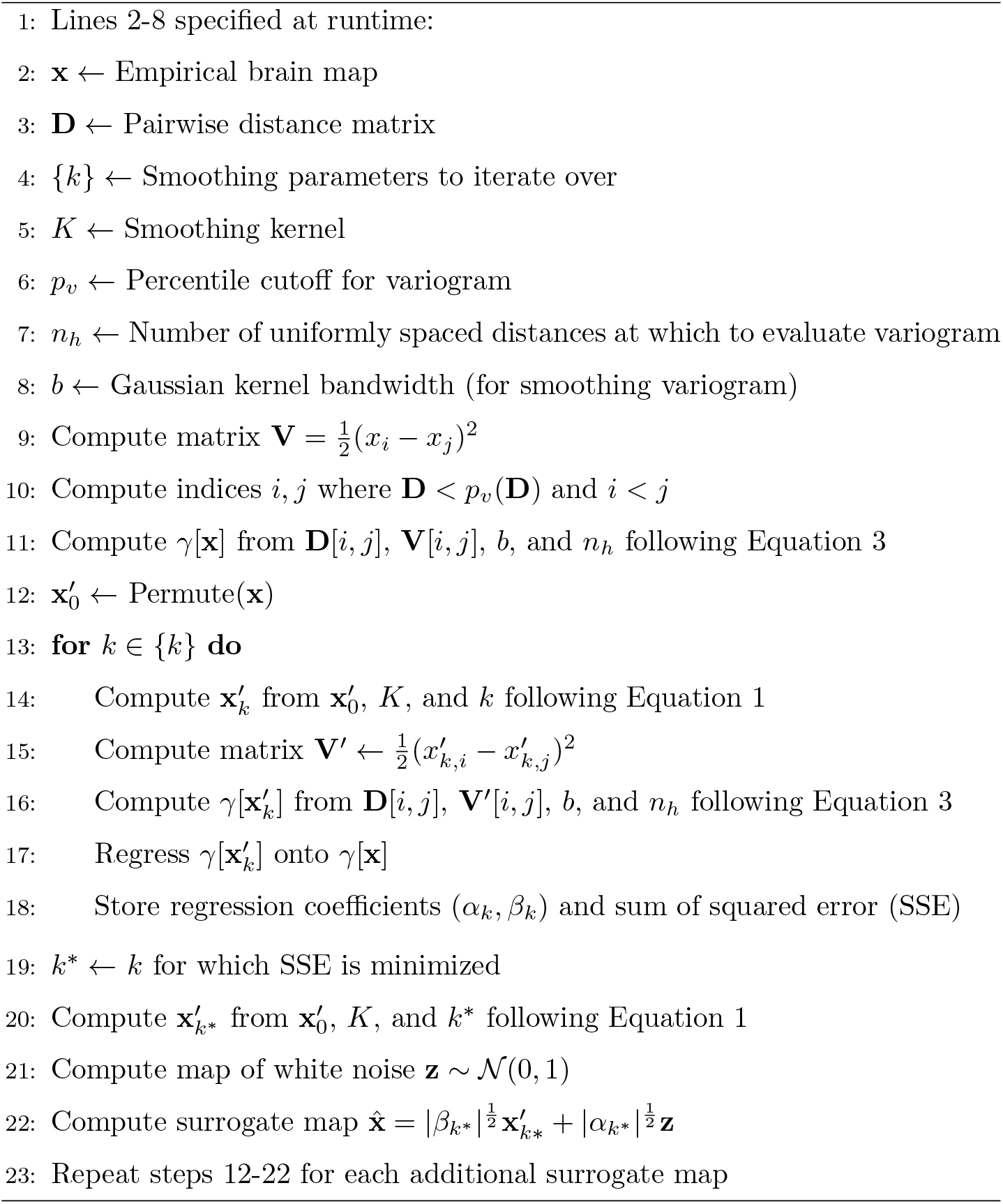

### Algorithm 2 Generating a dense SA-preserving surrogate map

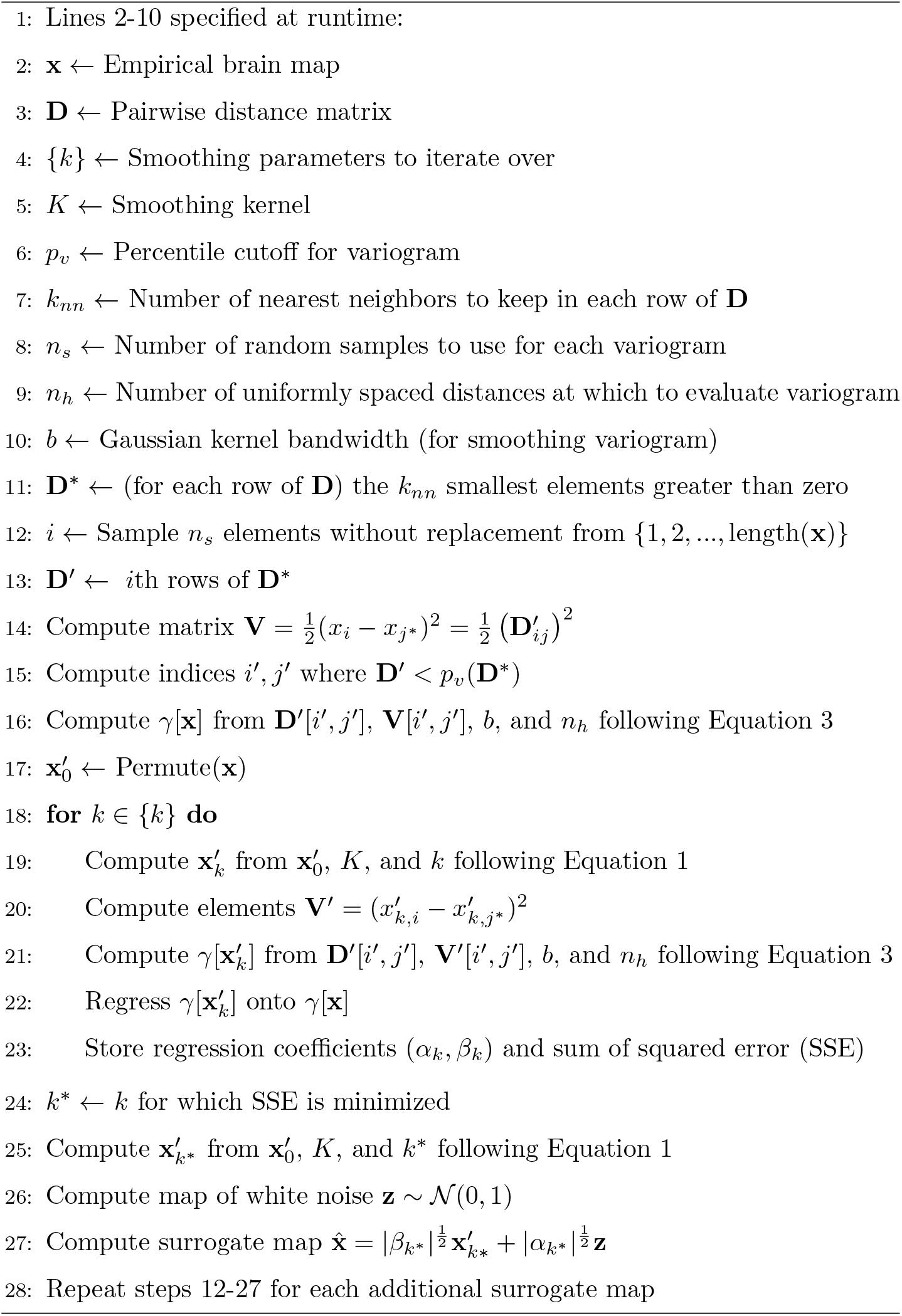

## Appendix B. Gaussian random fields

The most important statistical feature of a random field is its two-point autocorrelation function, *ξ*(*x*_1_, *x*_2_), which quantifies the inter-dependence of field values at different locations. For isotropic and homogeneous random fields, the autocorrelation function depends only on the separation *x* = *x*_1_ – *x*_2_ and is related to its power spectral density *P*(*k*) through a Fourier transform:

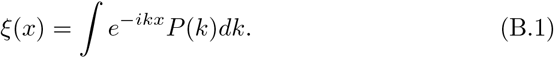

Because *ξ*(*x*) is positive definite, *P*(*k*) must be non-negative. In addition, for the field to be isotropic we require *P*(*k*) = *P*(|*k*|). In this spectral representation, field fluctuations are represented as an integral over plane waves, where *P*(*k*) specifies the distribution over wave number *k* (i.e., over spatial frequencies). In other words, the smoothness of the field is determined by the rate at which *P*(*k*) approaches 0 as *k* → ∞.

To simulate GRFs, we color a continuous white noise Gaussian process with zero mean and unit variance, *η*(*x*), whose Fourier transform is given by:

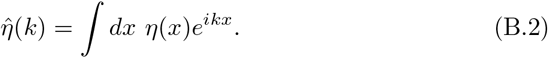

The Gaussian white noise process is then colored using the target power spectral density via:

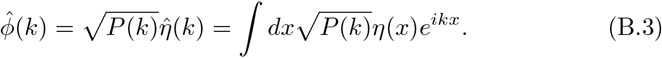

We then leverage the fact that the inverse Fourier transform of Equation B.3 yields a Gaussian process *φ*(*x*) colored with the desired power spectral density *P*(*k*) (Yura and Hanson, 2011). This field is guaranteed to be Gaussian by the central limit theorem because field values in *η*(*x*) are by definition independent, and an integral is an infinite sum. Note however that in discretized simulations where integrals are replaced by finite sums, this guarantee holds only asymptotically. Lastly, we let

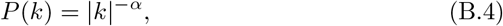

where *α* is any positive number, such that both the isotropic and positive-definiteness conditions on *ξ*(*x*) are satisfied.

To simulate GRF realizations, we follow the logic described above. We first create a uniformly spaced two-dimensional lattice (or grid) with *N* tilings in each dimension. At each point on the grid, we draw a random sample from a standard normal distribution. We then compute the discrete Fourier transform of this field to get 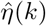. At each point on the grid, the shifted Fourier components 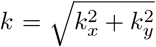 are used to compute the spectral amplitudes 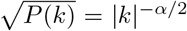. Note that we remove the mean shift by setting the spectral amplitude for *k* = 0 equal to zero. Finally, we substitute 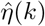 and 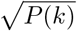 into Equation B.3 to get 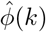, then take the discrete inverse Fourier transform to get *φ*(*x*). We normalize the resulting field such that it has zero mean and unit variance.

---

**Supplementary Figure 1:**
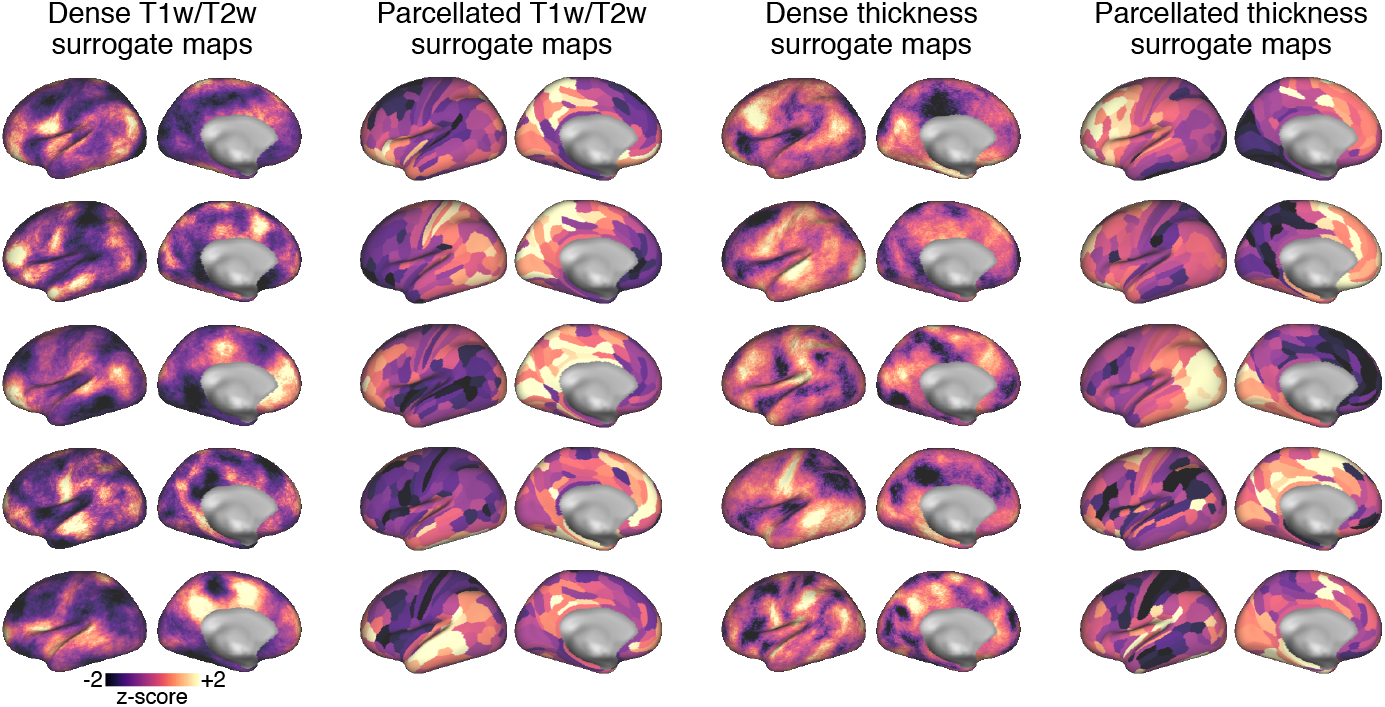
Random realizations of SA-preserving surrogate maps for the dense and parcellated group-averaged (*N* = 339) cortical T1w/T2w and cortical thickness maps. Each surrogate map’s topography is randomized while preserving the SA that is present in the empirical map from which each surrogate is derived. To match surrogate map value distributions to the distribution of values in the corresponding empirical map, rank-ordered surrogate map values were re-assigned the corresponding rank-ordered values in the empirical map.

**Supplementary Figure 2:**
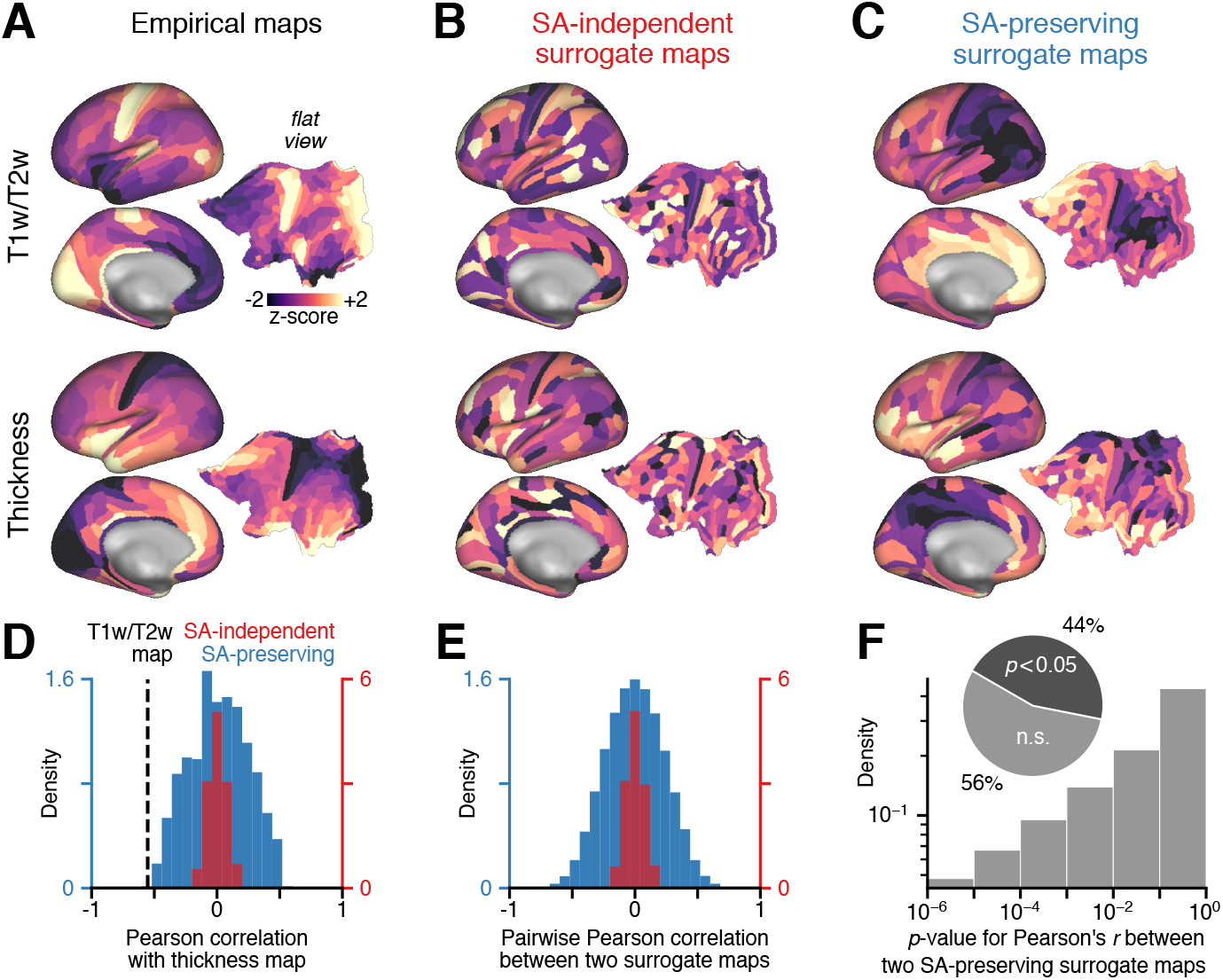
SA in parcellated brain maps has a substantial impact on statistical inference. (**A**) The group-averaged (*N* = 339) cortical T1w/T2w (top) and cortical thickness (bottom) maps across 180 parcels (Glasser et al., 2016a) in the left hemisphere. (**B**) Randomly shuffling the empirical maps in A destroys their autocorrelation structure. (**C**) One example realization of SA-preserving surrogate maps, derived for each empirical map. (**D**) Null distributions of Pearson correlation coefficients between the empirical cortical thickness map, and *N* = 1,000 randomly shuffled (red) and SA-preserving (blue) surrogate maps derived from the empirical T1w/T2w map. Dashed black line indicates the empirical correlation between the T1w/T2w and cortical thickness maps. (**E**) Distributions of Pearson correlation coefficients between pairs of randomly shuffled (red) and SA-preserving (blue) surrogate maps derived from the empirical T1w/T2w map. (**F**) Distributions of naive *p*-values for the Pearson correlation between pairs of SA-preserving surrogate maps derived from the empirical T1w/T2w map. These naive *p*-values are derived under the assumption that samples are independent and normally distributed, in which case the sampling distribution of Pearson’s *r* is a *t*-distribution with *n* – 2 = 178 degrees of freedom. n.s.: not significant.

**Supplementary Figure 3:**
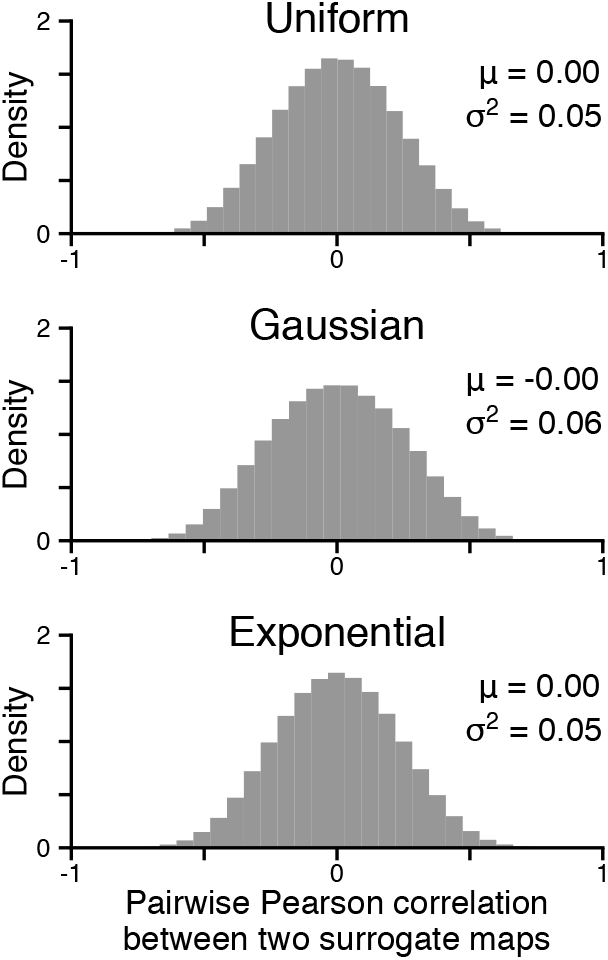
The SA-preserving surrogate generating method is largely insensitive to the functional form of the smoothing kernel (Equation 1). Each distribution consists of all pairwise Pearson correlations between *N* =1,000 SA-preserving surrogate maps derived from the parcellated cortical T1w/T2w map. Top, middle, and bottom panels were derived using uniform (i.e., distance independent), Gaussian, and exponentially-decaying kernels, respectively. Mean and variance of each distribution is indicated by *μ* and *σ*^2^, respectively.

**Supplementary Figure 4:**
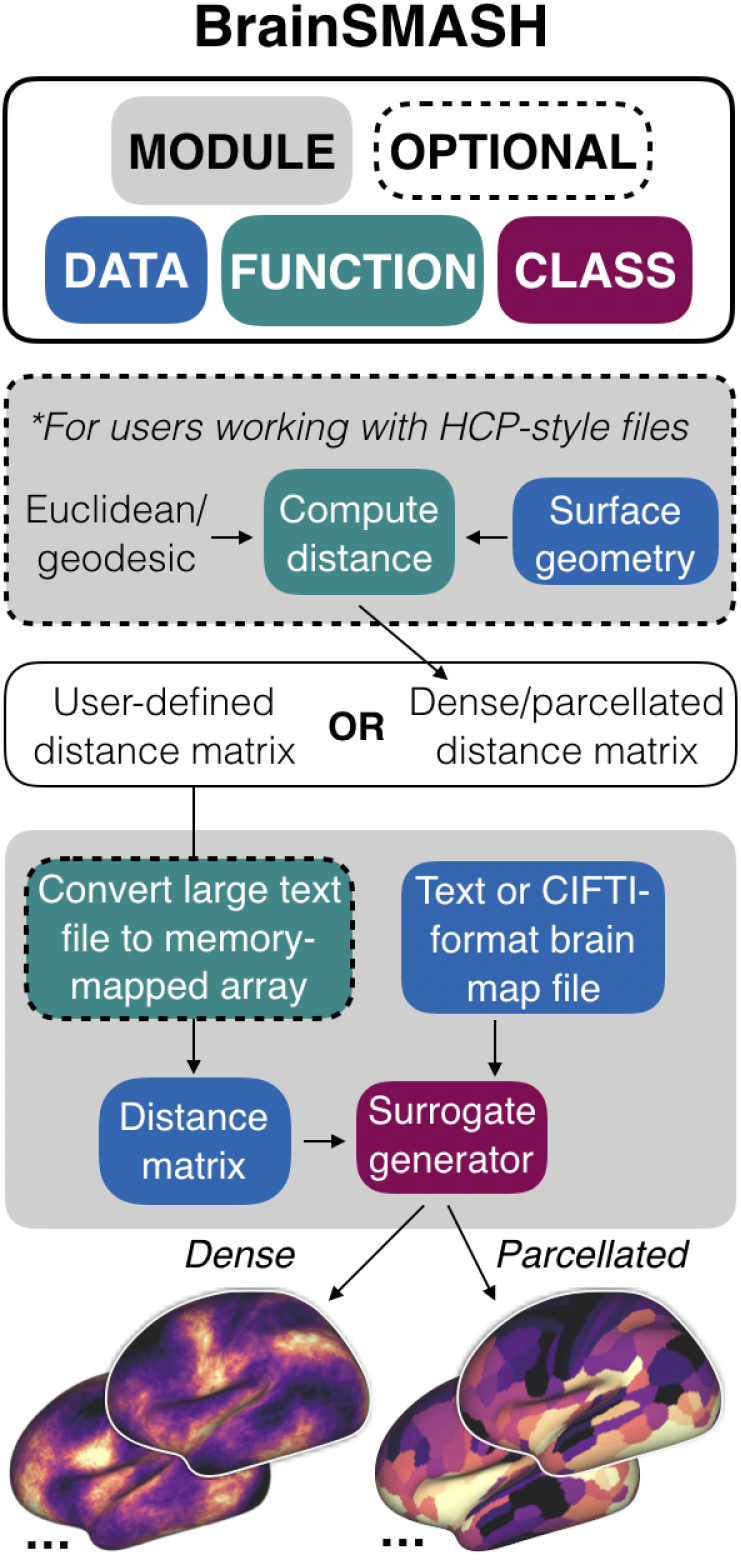
BrainSMASH: Brain Surrogate Maps with Autocorrelated Spatial Heterogeneity. Simulating surrogate brain maps with the BrainSMASH package requires specifying two inputs: a brain map, and the matrix of pairwise distances between elements of the brain map. Additional support is provided for users working with HCP-style neuroimaging files. Comprehensive documentation for BrainSMASH can be found at https://brainsmash.readthedocs.io.

**Supplementary Figure 5:**
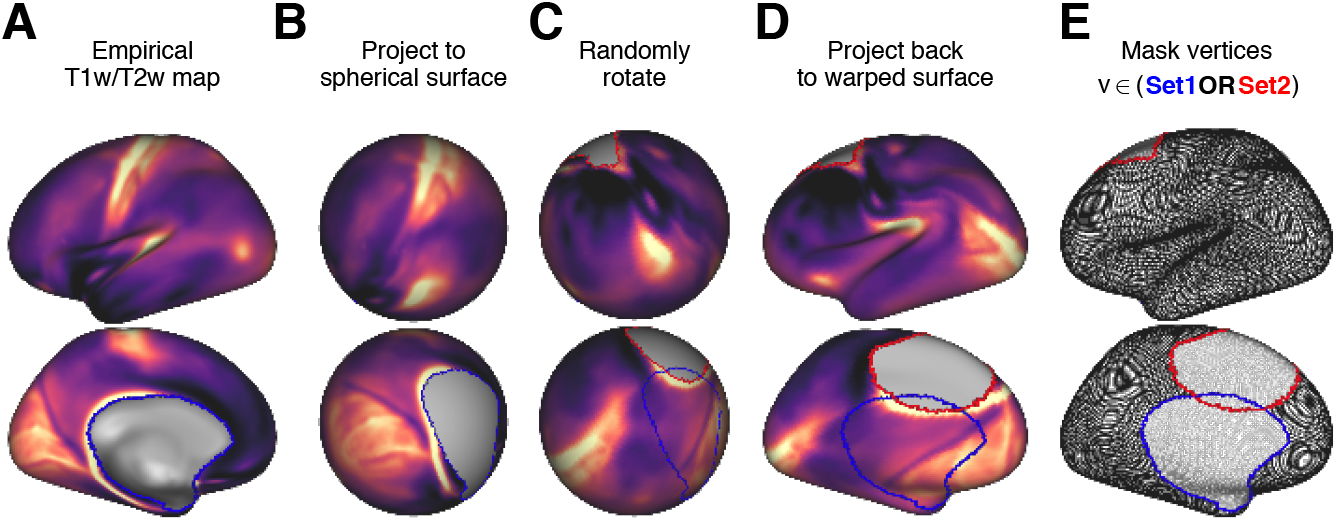
The spin test developed by Alexander-Bloch et al. (2018). (**A**) The dense, left-hemispheric cortical T1w/T2w map. The medial wall (blue border) forms a void in the cortical surface. (**B**) First, data are projected from the warped cortical surface onto a spherical surface. (**C**) Next, data on the spherical surface are rotated by a random angle, preserving SA but randomizing anatomical alignment. Blue (red) border indicates the location of the medial wall before (after) rotation. (**D**) Next, the rotated data are projected back onto the warped cortical surface. (**E**) To construct one sample from the null distribution, the rotated map in **D** is compared against another empirical brain map. Vertices in the pre- or post-rotation medial wall, indicated by lightly shaded edges within the blue and red borders, are excluded. The number of vertices included, indicated by darkly shaded edges, depends the spatial overlap between the pre- and post-rotation medial wall, and will in general vary for each sample in the null distribution. If a surrogate brain map were to be constructed, data rotated into the blue region would be discarded, while the red region would need to be interpolated in some manner using data points around the boundary.

**Supplementary Figure 6:**
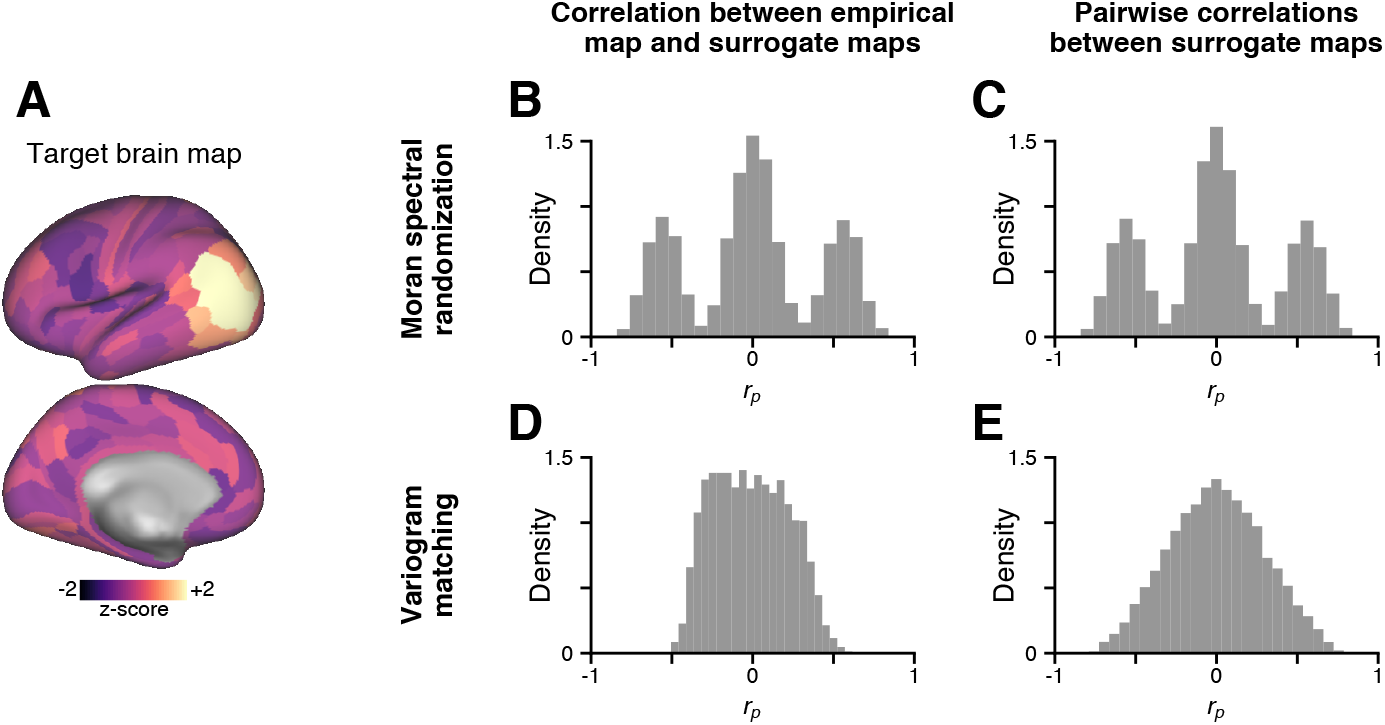
Moran spectral randomization (MSR) may yield surrogate brain maps which correlate strongly with the target map. (**A**) An example, neurobiologically plausible target brain map, constructed by superimposing normally distributed noise with a signal component which decays smoothly as a function of distance from area MT. (**B**) The distribution of Pearson correlations between the target map and *N* = 5, 000 MSR-derived surrogate maps, constructed using the BrainSpace toolbox (de Wael et al., 2019). A substantial fraction of MSR-derived surrogate maps correlate strongly (positively or negatively) with the target map, producing the two approximately symmetric outer peaks of the distribution. (**C**) The distribution of Pearson correlations between pairs of MSR-derived surrogate maps exhibits the same trimodal shape. Panels **B-C** indicate that MSR does not generate surrogate brain maps with random topographies. (**D**) The distribution of Pearson correlations between the empirical map and *N* = 5, 000 variogram-matched surrogate maps from our method. (**E**) The distribution of Pearson correlations between pairs of variogram-matched surrogate maps from our method.

## Notes

https://github.com/murraylab/brainsmash

https://brainsmash.readthedocs.io/

